# Functional interaction of low-homology FRPs from different cyanobacteria with *Synechocystis* OCP

**DOI:** 10.1101/247882

**Authors:** Yury B. Slonimskiy, Eugene G. Maksimov, Evgeny P. Lukashev, Marcus Moldenhauer, Cy M. Jeffries, Dmitri I. Svergun, Thomas Friedrich, Nikolai N. Sluchanko

**Author notes:** these authors equally contributed to the study (Yury B. Slonimskiy, Eugene G. Maksimov). Correspondence to: Nikolai N. Sluchanko. List of abbreviations: OCP: carotenoid protein, holoprotein OCP^AA^: OCP with amino acid substitutions Y201A and W288A, holoprotein NTE: N-terminal extension (comprising the αA helix up to amino acid 20) ∆NTE: OCP with the 12 most N-terminal amino acids deleted, holoprotein CTD: C-terminal domain NTD: N-terminal domain HCP: helical carotenoid protein, NTD homologue of OCP COCP: CTD of *Synechocystis* OCP (amino acids 165–317), holoprotein RCP: red carotenoid protein, holoprotein RCP(apo): NTD of *Synechocystis* OCP (amino acids 1–164), apoprotein FRP: fluorescence recovery protein SynFRP: *Synechocystis* FRP AnaFRP: *Anabaena* FRP AmaxFRP: *Arthrospira* FRP *Arthrospira*: *Arthrospira maxima* CS-328 *Anabaena*: *Anabaena variabilis* PCC 7937 *Synechocystis*: *Synechocystis sp.* PCC 6803 CAN: canthaxanthin ECN: echinenone hECN: 3’-hydroxyechinenone AL: actinic light LED: light-emitting diode DLS: dynamic light scattering DTT: dithiothreitol PB: phycobilisome QELS: quasi-elastic light scattering RC: reaction center of photosystem ROS: reactive oxygen species SEC: size-exclusion chromatography SEC-MALLS: SEC with multiangle laser light scattering analysis SAXS: small-angle X-ray scattering SDS-PAGE: sodium dodecyl sulfate polyacrylamide gel electrophoresis MSA: multiple sequence alignment UV: ultraviolet.

## Abstract

Photosynthesis requires a balance between efficient light harvesting and protection against photodamage. The cyanobacterial photoprotection system uniquely relies on the functioning of the photoactive orange carotenoid protein (OCP) that under intense illumination provides fluorescence quenching of the light-harvesting antenna complexes, phycobilisomes. The recently identified fluorescence recovery protein (FRP) binds to the photoactivated OCP and accelerates its relaxation into the basal form, completing the regulatory circle. The molecular mechanism of FRP functioning is largely controversial. Moreover, since the available knowledge has mainly been gained from studying *Synechocystis* proteins, the cross-species conservation of the FRP mechanism remains unexplored. Besides phylogenetic analysis, we performed a detailed structural-functional analysis of two selected low-homology FRPs by comparing them with *Synechocystis* FRP (*Syn*FRP). While adopting similar dimeric conformations in solution and preserving binding preferences of *Syn*FRP toward various OCP variants, the low-homology FRPs demonstrated distinct binding stoichiometries and differentially accentuated features of this functional interaction. By providing clues to understand the FRP mechanism universally, our results also establish foundations for upcoming structural investigations necessary to elucidate the FRP-dependent regulatory mechanism.

## Introduction

Due to the well-known threats of reactive oxygen species (ROS), all photosynthetic organisms are forced to balance between photosynthesis and photoprotection (Peschek, 2011). Carotenoids are critical in mediating this process as they avert the accumulation of ROS (Pascal et al, 2005). Carotenoids can compete with photosynthetic reaction centers (RCs) for excitation energy and effectively dissipate the absorbed energy excess into heat thus allowing plants, algae and cyanobacteria to adapt to different environmental conditions. In cyanobacteria, the presence of water-soluble extramembrane antenna complexes called phycobilisomes (PBs) – which are substantially different from intramembrane light-harvesting complexes of plants (Adir, 2005) – necessitates the coupling with a specific type of water-soluble carotenoid-binding protein, the Orange Carotenoid Protein (OCP). The first OCP was purified *inter alia* from *Arthrospira maxima* in 1981 (Holt & Krogmann, 1981) and the genetic sequence determined in 1997 (Wu & Krogmann, 1997) while the atomic structure was solved in 2003 (Kerfeld et al, 2003), i.e., long before the functional role was fully established (Karapetyan, 2007; Wilson et al, 2006).

OCP is a molecular photoswitch that upon absorbing a blue-green photon (420–550 nm) undergoes a spectral red shift from the basal, dark-adapted orange state, OCP^O^, to the red-shifted, metastable quenching state, OCP^R^. The key to phototransformation is the light absorption by a single keto-carotenoid chromophore (in OCPs from native sources 3’-hydroxyechinenone, hECN) that triggers significant rearrangements of the 35 kDa protein matrix (Wilson et al, 2008). The quantum yield of this process is about 0.2 % (Maksimov et al, 2015; Maksimov et al, 2017c) that is sufficient for keeping OCP in its inactive orange form under low to moderate insolation levels suitable for photosynthesis. The stability of the orange form is determined by multiple protein-chromophore interactions and structural features of the protein matrix. The structural characteristics of OCP are dominated by two structurally distinct N- and C-terminal domains (NTD and CTD, respectively) in addition to: (i) a flexible interdomain linker; (ii) an N-terminal extension (NTE) that interacts with a specific site in the C-terminal domain; (iii) numerous contacts between the domains in the carotenoid-binding cavity (Bandara et al, 2017; Liu et al, 2014; Sluchanko et al, 2017c; Thurotte et al, 2015; Wilson et al, 2012; Wilson et al, 2011); and (iv) two distinct H-bonds between a Trp and a Tyr residue in the CTD and the keto-oxygen of the chromophore. The formation of the active form is accompanied by reversible disruption of a majority of the interactions listed above, a 12 Å translocation of carotenoid into the N-terminal domain, and a complete separation of the NTD and CTD (Gupta et al, 2015; Leverenz et al, 2015; Maksimov et al, 2017a). These molecular events hinder the light-independent back reaction and allows OCP to adopt the active form long enough to bind to the PBs core and quench PBs fluorescence.

An active PBs quenching form of OCP may also be obtained by the destabilization of protein-chromophore interactions due to mutation of key Tyr-201/Trp-288 residues (Maksimov et al, 2016; Maksimov et al, 2017c; Sluchanko et al, 2017a). Additionally, PBs fluorescence can be quenched by the isolated carotenoid-containing NTD formed upon partial proteolysis of OCP (designated red carotenoid protein, RCP) (Leverenz et al, 2014). These observations have raised the idea about the functional modularity of OCP and have stimulated research of the individual properties of the CTD, NTD and isolated homologues thereof, the genes of which are present in different families of cyanobacteria along with full-length OCP genes (Lechno-Yossef et al, 2017; Lopez-Igual et al, 2016; Maksimov et al, 2017b; Melnicki et al, 2016; Moldenhauer et al, 2017a; Muzzopappa et al, 2017).

OCP-mediated PBs fluorescence quenching is controlled by another water-soluble factor, the Fluorescence Recovery Protein (FRP) (Boulay et al, 2010; Gwizdala et al, 2013; Sutter et al, 2013). *In vitro,* FRP significantly increases the rate of OCP^R^ relaxation to OCP^O^ (Boulay et al, 2010; Sluchanko et al, 2017a; Sluchanko et al, 2017c; Sutter et al, 2013) and destabilizes OCP^R^-PBs complexes, which results in restoration of full antenna capacity (Gwizdala et al, 2011; Thurotte et al, 2017). However, the molecular mechanism of FRP binding to OCP is largely unknown. The main site of FRP-OCP interaction is thought to be located in the CTD, which is supported by the ability of FRP to bind to several OCP forms with separated domains (Sluchanko et al, 2017a), to the individual CTD (Moldenhauer et al, 2017a; Sutter et al, 2013), and also to the ΔNTE mutant with non-separated domains but with exposed tentative FRP-binding site(s) (Sluchanko et al, 2017c). It was found that, while normally forming stable dimers (Lu et al, 2017; Sluchanko et al, 2017a), after binding to OCP, FRP can undergo monomerization (Moldenhauer et al, 2017b; Sluchanko et al, 2017a; Sluchanko et al, 2017c), although the reason for and necessity of this dissociation is completely unclear. It was shown that FRP assists in the correct positioning of the CTD and NTD to facilitate carotenoid back translocation into the CTD and to accelerate the reformation of basal OCP^O^ (Maksimov et al, 2017c). Nevertheless, the structures of FRP complexes with various OCP forms, which would substantially clarify the FRP action mechanism, are unknown.

Like OCP, FRP homologues are present in multiple different families of cyanobacteria (Bao et al, 2017). FRP amino acid sequences are typically far less than 50 % identical, whereas the primary structure of OCPs is much more conserved, usually above 80 %. This fact raises the principal question as to whether the FRP-mediated regulatory mechanism is universal across cyanobacteria or only species-specific because of mutual evolutionary adaptation of the interaction interfaces between OCP and FRP. To address this question, we compared the structures and functional activities of previously uncharacterized FRPs from *Anabaena variabilis* and *Arthrospira maxima,* identified as having only 38 % amino acid sequence identity with FRP from *Synechocystis sp.* PCC6803 (termed *Synechocystis* herein). Although significant differences in the amino acid sequences are present between these cyanobacterium species, we show that FRPs assemble into a rather conserved dimeric structure that adopts similar conformations in solution. Of interest, low-homology FRPs were able to interact with the well-described *Synechocystis* OCP and regulate OCP-induced non-photochemical quenching of PBs fluorescence, suggesting a common structure-based mode of the FRP-OCP regulation across species.

## Results

### Isolation and characterization of selected FRP homologues from different species

In contrast to OCP homologues, FRPs from different cyanobacteria are substantially more dissimilar and less well studied. Indeed, FRP was discovered only about seven years ago, and until very recently (Boulay et al, 2010, the only crystal structure available was that from *Synechocystis* (*Syn*FRP), showing two protein conformations assembled into dimeric and tetrameric forms (PDB 4JDX) (Sutter et al, 2013). The physiological importance of the tetrameric form is still controversial and several studies reported dimers as a prevalent oligomeric FRP form in solution (Lu et al, 2017; Sluchanko et al, 2017a). The dimeric assembly was recently observed in a novel crystallographic structure of FRP from *Fremyella diplosiphon* (*Tolypothrix* sp. PCC7601) (Bao et al, 2017), resembling that of *Syn*FRP with C_α_ root mean square deviation (rmsd) of 1.45 and 1.82 Å (depending on which chains are aligned). Despite the apparent structural similarity of the two FRP homologues, the universality of the FRP mechanism remained an unresolved question.

To directly address this question, we decided to study FRP homologues from different species having significantly dissimilar amino acid sequences compared to the well-characterized *Syn*FRP. On the basis of a bioinformatics analysis of fifty non-redundant FRP-like protein sequences (see Supplementary text 1 and Fig. S1), we built a phylogenetic tree showing the relationships between FRP homologues (Fig. 1A). For this study we selected representative FRP variants from *A. variabilis* (*Ana*FRP; Uniprot Q3M6D9) and *A. maxima* (*Amax*FRP; Uniprot B5W3T4), designated in the Uniprot database as “uncharacterized proteins” (and presented as several Uniprot entries each; see Materials and Methods), and produced them recombinantly in *Escherichia coli*. The multiple sequence alignment (MSA) of these variants and *Syn*FRP (Uniprot P74103) revealed only 37.6% identity among the three amino acid sequences (Fig. 1B).

**Fig. 1.**
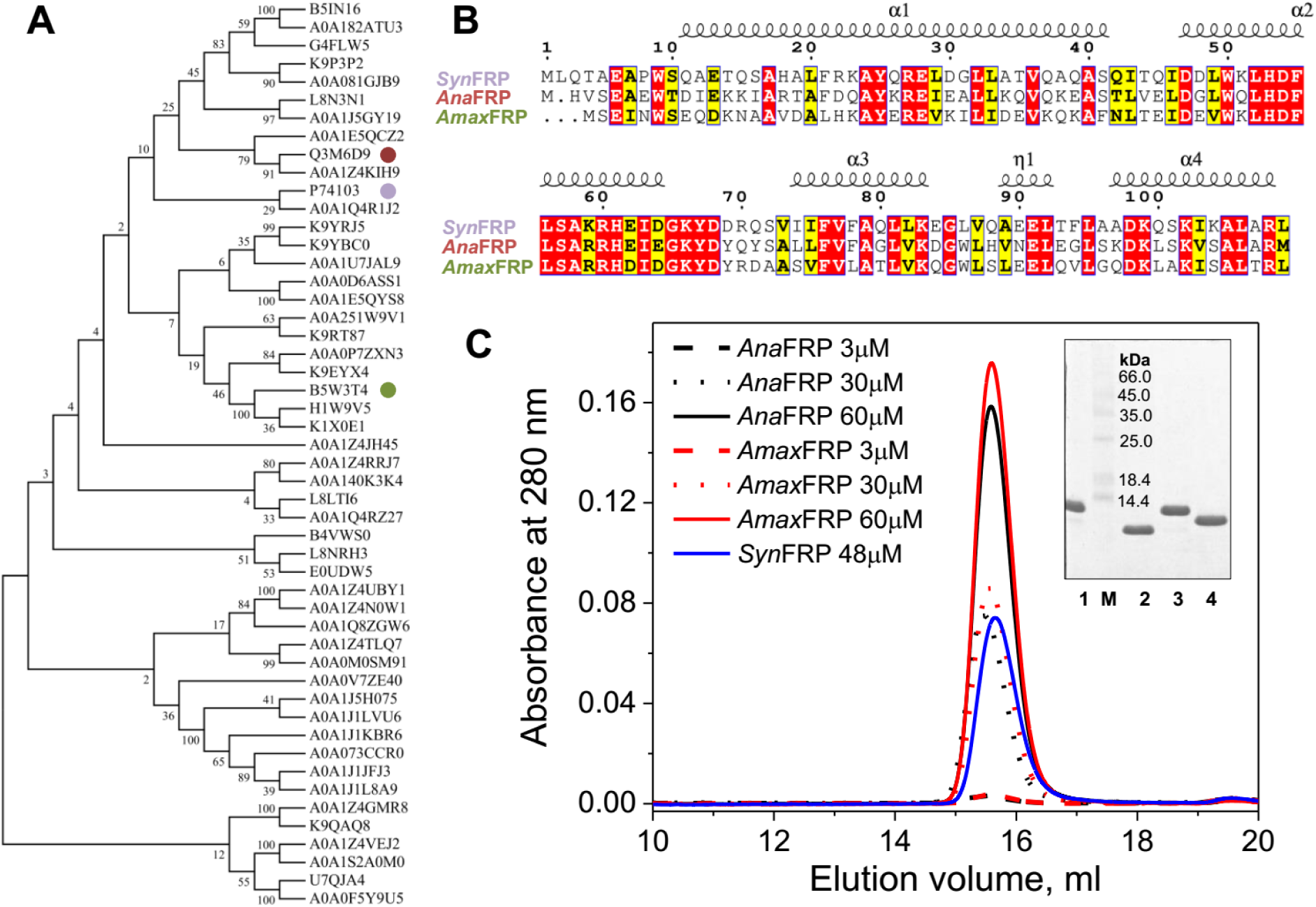
Characterization of FRP homologues from *Anabaena variabilis* and *Arthrospira maxima*. **A**. Phylogenetic consensus tree obtained for fifty FRP sequences (see Supplementary text 1) by the Maximum Likelihood method and consequent bootstrapping (Felsenstein, 1985; Jones et al, 1992; Kumar et al, 2016). The percentage of replicate trees in which the associated taxa clustered together in a bootstrap test (100 replicates) is shown next to the branches. Color coded circles mark three FRP species selected for the present study. **B.** MSA of *Synechocystis*, *Anabaena* and *Arthrospira* FRP homologues, characterized in this study, showing assignment of the secondary structure with colouring scheme considering physico-chemical similarity of amino acid residues. Identical residues are highlighted in red, similar ones in yellow. **C**. Concentration dependencies of the oligomeric state of *Ana*FRP and *Amax*FRP analyzed using a Superdex 200 10/300 column at a 1.2 ml/min flow rate. Concentrations of the protein samples loaded (100 μl) are indicated. Insert: purity of the *Syn*FRP (1), *Syn*FRP_8-109_ (2), *Ana*FRP (3), *Amax*FRP (4) preparations controlled by 17 % SDS-PAGE. M – protein markers with *M*_W_indicated to the right.

The purified FRP proteins were homogenous (Fig. 1C, insert) and demonstrated highly symmetrical peaks on size-exclusion chromatography (SEC), with positions almost unchanged upon 20-fold dilution and very similar to that of *Syn*FRP (Fig. 1C). Note that due to the substantial differences in extinction coefficients at 280 nm (due to different tryptophan content of the proteins), the amplitudes of the peaks of *Ana*FRP and *Amax*FRP at 30 μM were close to that of *Syn*FRP at 48 μM load concentration. Given the very similar behavior of *Ana*FRP and *Amax*FRP, only *Amax*FRP was selected for further structural analysis.

In order to characterize the structural conformation of the FRP proteins in solution we used small-angle X-ray scattering (SAXS). The initial wild-type *Syn*FRP construct described in our previous work (Sluchanko et al, 2017a) contained an uncleavable N-terminal His_6_ tag and a linker, making it significantly longer than the resolvable amino acids in the existing crystal structure (PDB 4JDX; residues 8–109), potentially complicating structural analyses. Hence, a truncated version of *Syn*FRP spanning amino acids 8–109 ( *Syn*FRP_8-109_) was engineered, with a calculated monomeric *M*_W_ of 11.6 kDa. The purified protein showed a concentration-dependent SEC elution profile spanning 4–200 μΜ load concentration (Fig. 2A), suggesting either oligomeric state transitions or conformational heterogeneity. Further analysis by SEC-MALLS/SAXS at a high-load protein concentration (460 μM) revealed a single symmetrical peak with a flat distribution of the *M*_W_ values determined from light scattering (Fig. 2B). The mean value of 28 kDa obtained from MALLS combined with the concentration-independent *M*_W_ estimates from the resulting SAXS profile suggested that *Syn*FRP_8-109_ forms dimers (Fig. 2C and Supplementary Table S1; *M*_W_ Porod = 23 kDa (Petoukhov et al, 2012); *M*_W_ volume-of-correlation = 25 kDa (Rambo & Tainer, 2013)). A comparison between the hydrodynamic radius, *R*_h_, obtained from DLS (2.86 nm) and the radius of gyration *R* _*g*_ from SAXS (2.91 nm) indicates that the shape factor, *R*_g_/ *R*_h_, of ~1 is much larger than that expected for globular/spherical particles ( *R*_g_/ *R*_h_=0.78). In combination with the skewed distribution real-space distances *p*( *r*) (Fig. S2) extending to a maximum size of *D* _*max*_ = 10.5 nm, this result indicates that dimeric *Syn*FRP_8-109_ adopts a highly extended structure. An *ab initio* shape model of the dimer directly generated from the SAXS data using *GASBOR* (discrepancy of the shape fit *χ*^2^ = 1.15) (Svergun et al, 2001) is shown in Fig. 2D spatially superimposed with the X-ray crystal structure (PDB 4JDX, chains A and C’). The extended conformation of the dimer is apparent in both models, however, unlike the *GASBOR* model, the scattering computed from the crystal structure does not fit the SAXS data well ( *χ*^2^=2.96, Fig. 2C). This discrepancy may originate from a shift in the angle of approach between the extended helical arms of opposing monomers that otherwise form the binding interface of the dimer. Indeed, the *ab initio* shape appears more ‘kinked’ compared to the crystal structure. A rigid-body refinement of the latter yielded a significant improvement in the fit to the SAXS data when allowing for a change in the angle between the two monomers of about 30° ( *χ*^2^=1.4; Fig. 2D). As a caveat, although the deletion of the very first residues in *Syn*FRP used in this study does not prevent dimer formation, we cannot disregard the possibility that the deletion may have slightly changed the conformation of the interface resulting in the sliding of the subunits, which at the same time may be a typical feature of conformational dynamics of various FRP proteins.

**Fig. 2.**
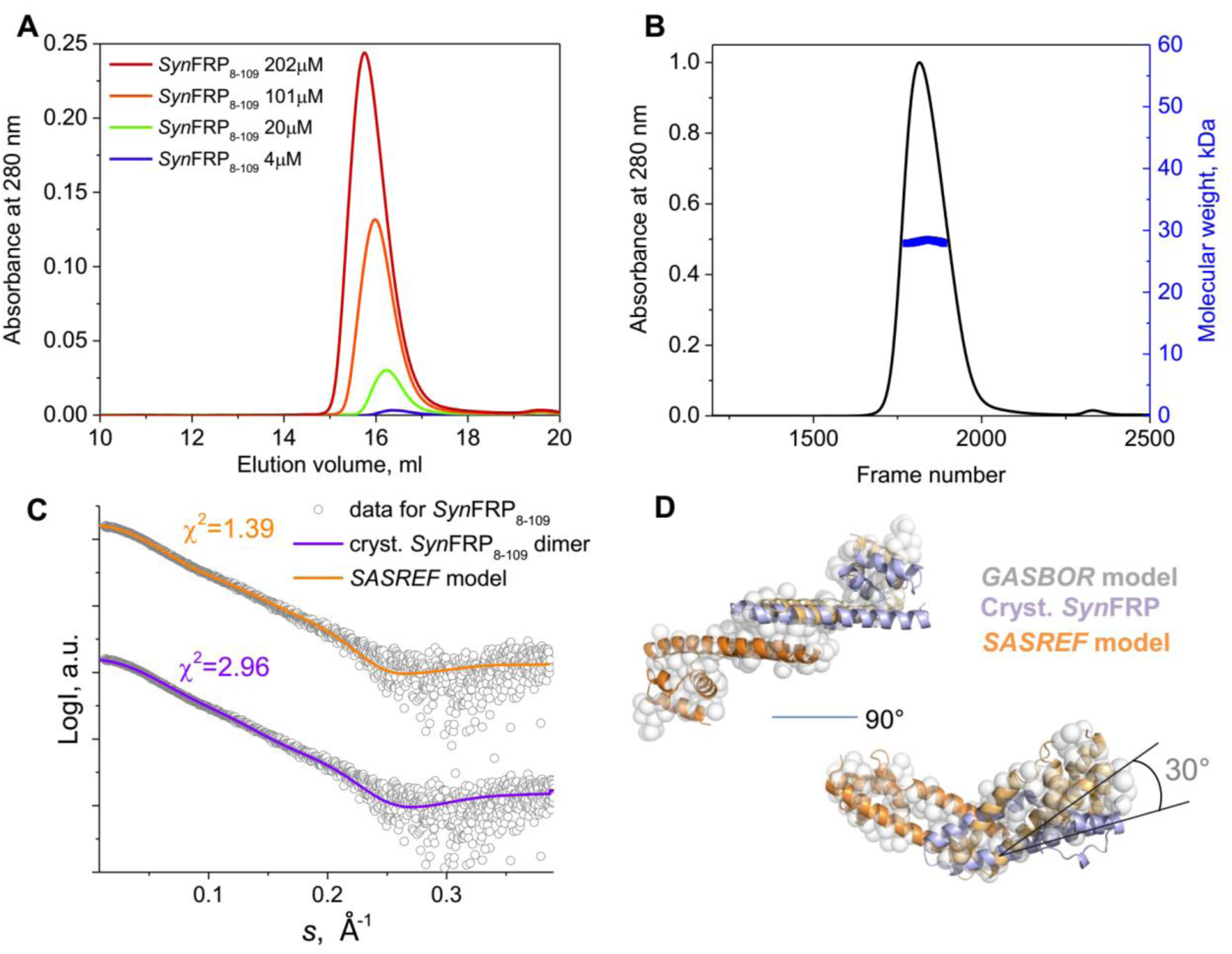
Hydrodynamic properties and solution conformation of *Syn*FRP_8-109_. **A**. SEC on a Superdex 200 Increase 10/300 column at four different load protein concentrations (indicated in μM). **B**. SEC-MALLS/SAXS analysis of *Syn*FRP_8-109_ showing the UV absorption trace and *M*_W_ distribution obtained from MALLS. The SAXS data were collected in parallel to MALLS. **C**. Fitting of the final SAXS profile obtained by averaging of the SAXS curves across the peak in **A** by the crystallographic *Syn*FRP dimer (PDB 4JDX, chains A and C’) or the dimer with the refined arrangement of the monomers as rigid bodies (program *SASREF*, see Methods). For clarity, the curves are shifted along the Y axis. **D**. Superposition of the *ab initio GASBOR* model (gray spheres), the crystallographic *Syn*FRP dimer (orange and violet subunits in ribbon representation) and the *SASREF* model thereof (orange and wheat subunits). The atomistic models were aligned by one monomer (orange) to reveal the angular shift.

An *Amax*FRP construct (monomer *M*_W_ = 12.6 kDa) was analyzed using batch SAXS experiments at different sample concentrations. No concentration-dependent effects were observed (data not shown). The data clearly indicate that the protein forms dimers in solution and that the overall structural parameters share common features with *Syn*FRP_8-109_. The experimental *M*_W_ obtained from different methods is very close to be twice the *M*_W_ of a monomer (Supplementary Table S1). The *R* _*g*_ (2.8 nm), *D* _*max*_ (9.5 nm) and the resulting skewed *p*( *r*) profile (Fig. S2) once again show that the dimers are structurally anisotropic as is also revealed by the *GASBOR* model (Fig. 3B). As there is no X-ray crystal structure available for *Amax*FRP we first assessed how well the FRP homologues from *Synechocystis* (PDB 4JDX, chains A and C’) and *Tolypothrix* (PDB 5TZ0) fit the scattering data. Although the fits appear reasonable, significant systematic discrepancies are present when comparing the model with experimental scattering intensities (Fig. 3A). To account for differences in the primary structures between the homologues, we built a homology model for *Amax*FRP using iTASSER (Yang et al, 2015); this model provided an excellent fit to the SAXS data ( *χ*^2^ = 1.13; Fig. 3A) and spatially aligns to the *ab initio GASBOR* model (Fig. 3B). The primary difference between the arrangement of *Amax*FRP and *Syn*FRP_8-109_ is the angle of approach at the interface between the extended helical arms of the monomers ( *Amax*FRP ~135°; *Syn*FRP_8-109_ ~105°).

**Fig. 3.**
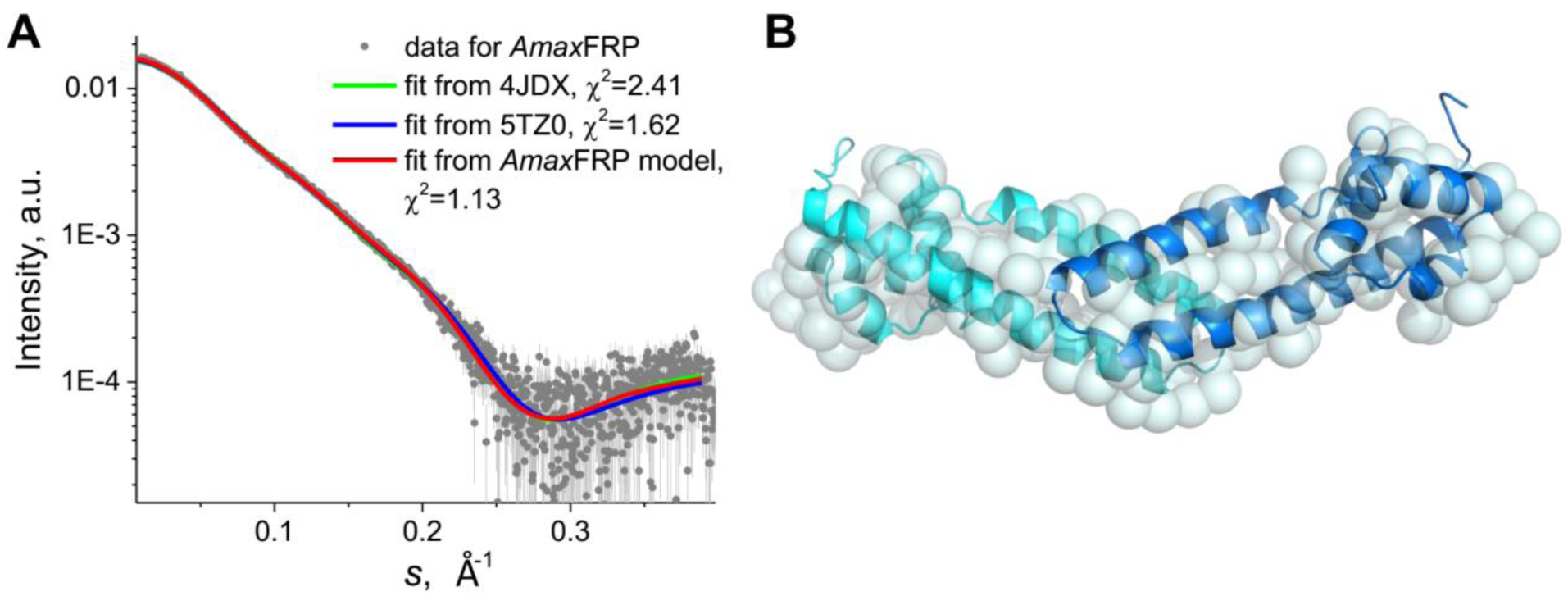
Solution conformation of *Amax*FRP studied by SAXS. **A**. Fitting of the SAXS curve collected for *Amax*FRP at 460 μM protein concentration by the atomistic homology model of the *Amax*FRP dimer obtained using *iTASSER* (Yang et al, 2015). The fit to the data for the X-ray crystal structures of *Synechocystis* and *Tolypothrix* FRP homologues (PDB 4JDX and 5TZ0) are also displayed. **B**. Overlay of the *Amax*FRP dimer (subunits in blue and cyan) and the best fitting *ab initio GASBOR* bead model (semitransparent cyan spheres).

Such a combined analysis allows us to hypothesize that FRP proteins form extended dimers with similar conformations that may differ in the angle between the helical arms of the monomers at the dimer-subunit interface. However, considering the wide diversity of FRP-like homologues (Fig. 1A), we cannot as yet predict whether such conformations are generally applicable across the entire FRP protein family.

### Direct binding of the FRP homologues to Synechocystis OCP and its derivatives

In order to understand whether dimeric FRPs with dissimilar amino acid sequences share a universal mechanism of binding to OCP and mutants or individual domains thereof, we analyzed the direct interaction of *Syn*FRP, *Ana*FRP, *Amax*FRP with *Synechocystis* OCP (Fig. 4A-C) and variants including: (i) the presumably strongest FRP binder, an OCP variant lacking the N-terminal extension (NTE) that was proposed to mask the FRP binding side in OCP^O^ (∆NTE; Fig. 4D-F and Fig. 5) (Sluchanko et al, 2017c); (ii) an analog of the active signaling OCP form with separated domains (OCP^AA^ mutant (Maksimov et al, 2017c; Sluchanko et al, 2017a); Fig. 4G-I) and; (iii) individual domains of OCP (Fig. 6). Our previous biochemical studies proved analytical SEC to be especially efficient for fast assessment of the presence of various protein-protein interactions involving OCP and its derivatives (Maksimov et al, 2017b; Sluchanko et al, 2017a; Sluchanko et al, 2017c).

**Fig. 4.**
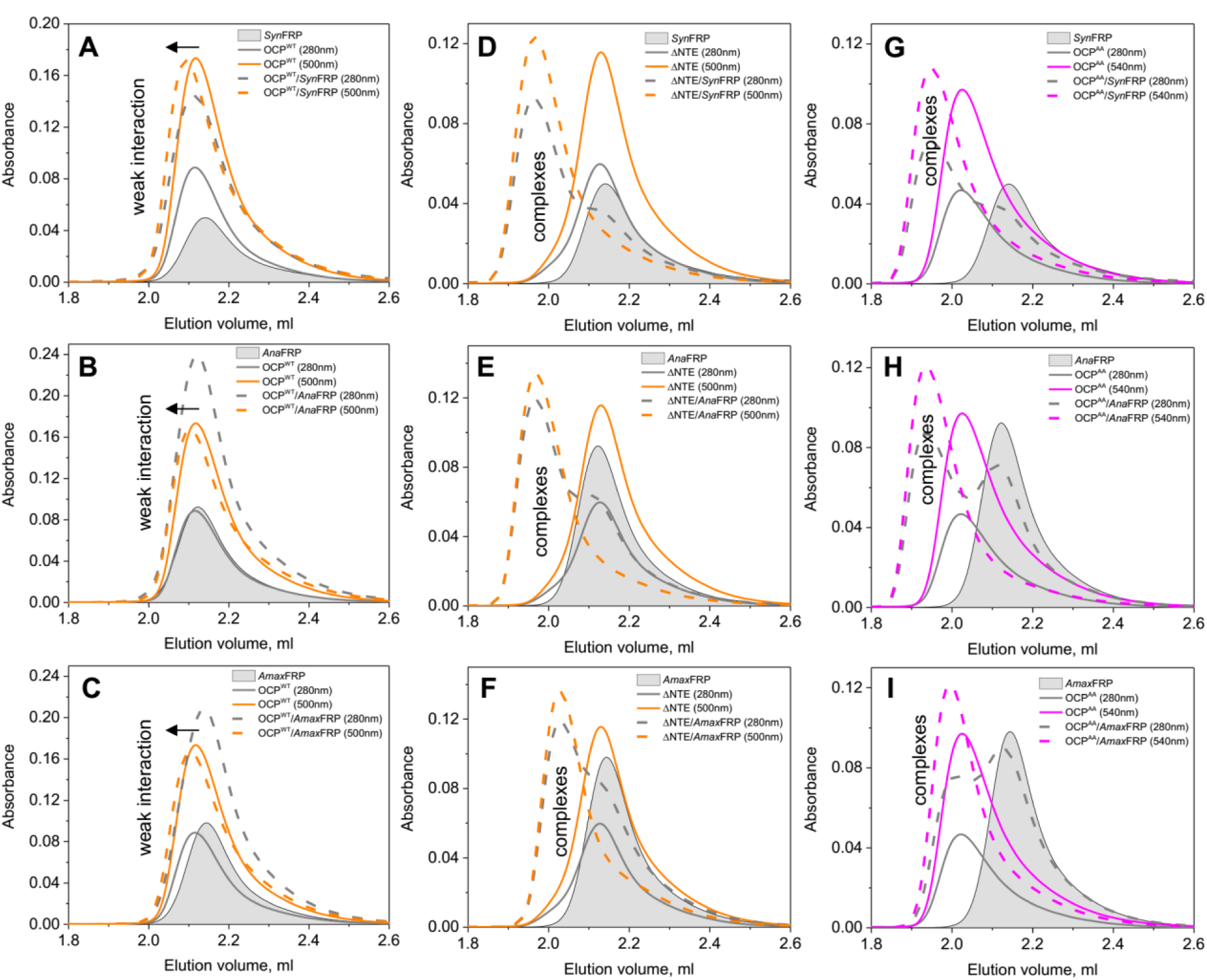
Interaction of *Syn*FRP (A, D, G), *Ana*FRP (B, E, H), or *Amax*FRP (C, F, I) with either OCP^WT^ (A-C), ΔNTE (D-F), or OCP^AA^ (G-I) analyzed by SEC. The samples (40 μl) containing individual OCP variants (10 μM, solid lines), FRP species (20 μM; semitransparent gray peaks) or mixtures of various OCP (10 μM) and FRP (20 μM) (dashed lines) were run on the pre-equilibrated Superdex 200 Increase 5/150 column followed by 280 nm (gray lines) and visible absorption (wavelengths are indicated; color of the lines roughly corresponds to that of samples) at a flow rate of 0.45 ml/min. Due to the low expected affinity to OCP^WT^ (Sluchanko et al, 2017a), higher concentrations of FRPs were used in A-C (40 μM instead of 20 μM in other cases). Arrows in A-C indicate the shift reflecting weak protein-protein interactions. Discernible complexes in D-I are marked.

**Fig. 5.**
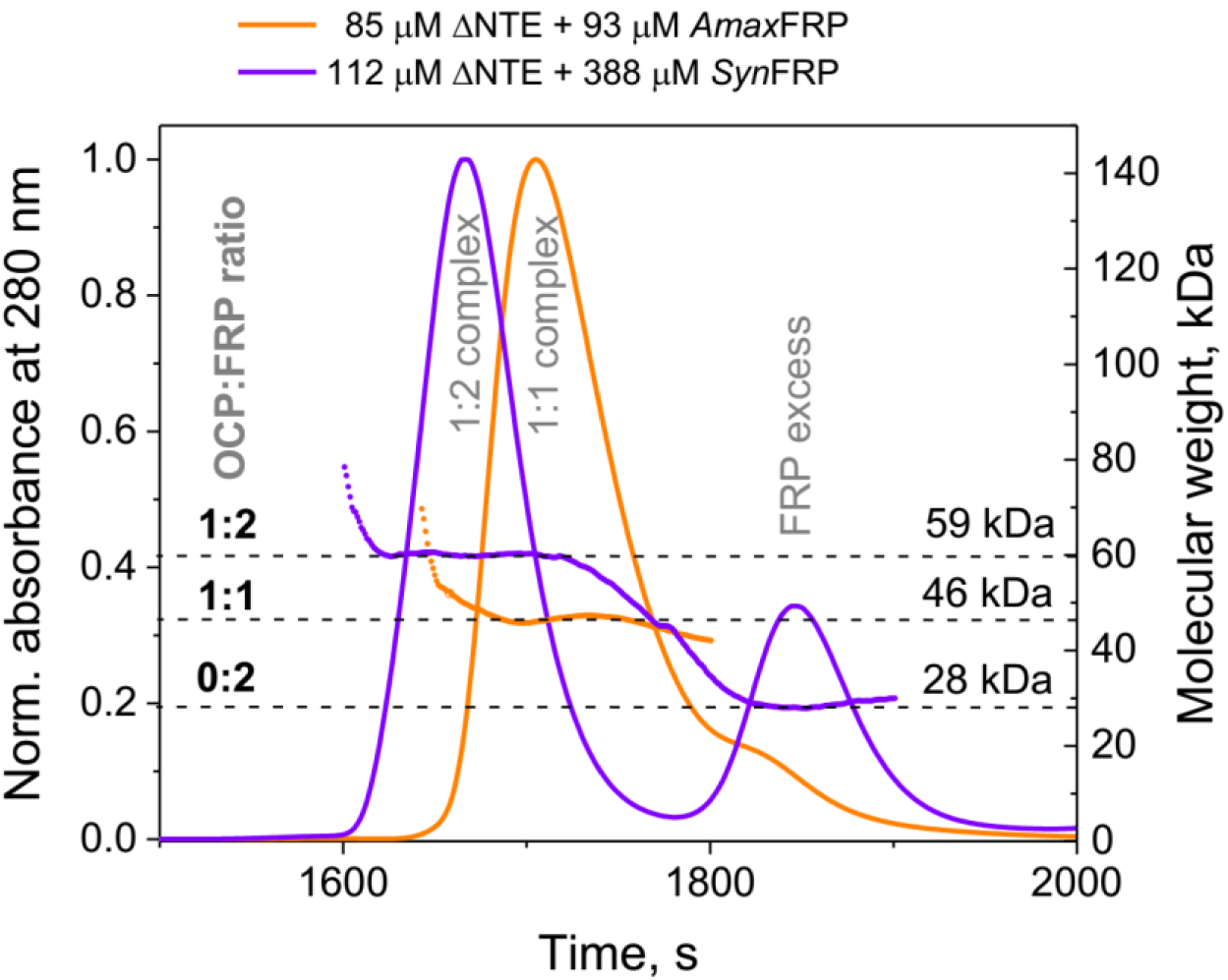
The ability of two different FRPs to form either 2:1 or 1:1 complexes with ΔNTE demonstrated by SEC-MALLS. Analysis of the ΔNTE mixtures with either *Syn*FRP or *Amax*FRP was performed at different molar FRP excess using a Superdex 200 Increase 10/300 column coupled with MALLS. Concentrations of the proteins in the pre-incubated mixtures (100 μl each) are indicated on top, the profiles are normalized to the maximum of the complex peak for clarity. Mean *M*_W_ values across the peaks of the 1:1 or 1:2 OCP complexes or *Syn*FRP excess, obtained from light scattering data, are shown by dashed lines. Flow rate: 0.5 ml/min. Temperature: 20 °C.

**Fig. 6.**
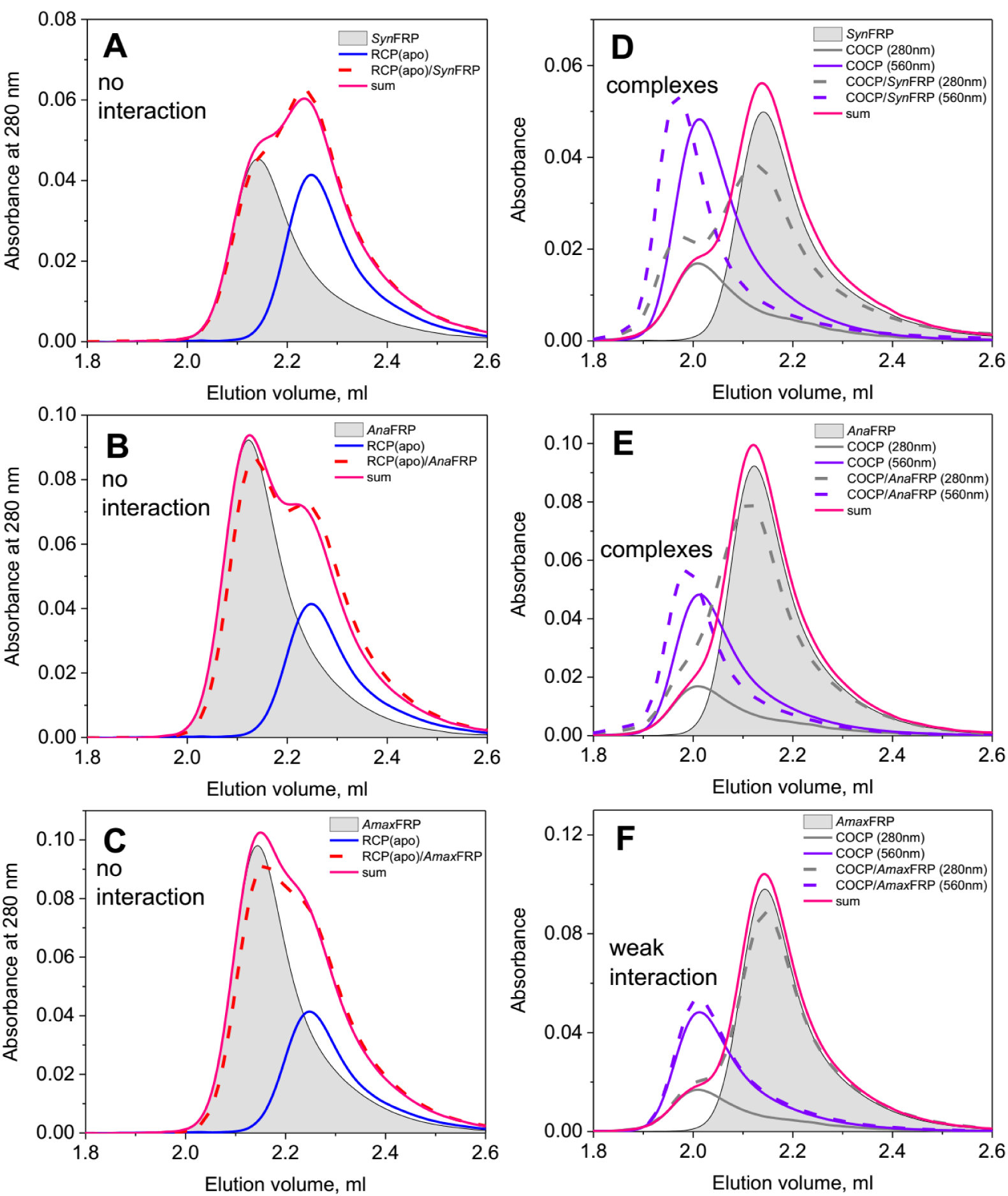
Assessment of interaction between either *Syn*FRP (A, D), *Ana*FRP (B, E), or *Amax*FRP (C, F) with OCP domains – RCP(apo) (A-C) or COCP (D-F) – analyzed by SEC. The samples (40 μl) containing individual OCP derivatives (10 μM), FRP species (20 μM; semitransparent gray peaks) or mixtures of OCP derivatives (10 μM) and FRPs (20 μM) (dashed lines) were run on the pre-equilibrated Superdex 200 Increase 5/150 column followed by 280 nm (gray lines) and 560 nm (only in the COCP case) at a flow rate of 0.45 ml/min. The algebraic sum of the individual 280-nm elution profiles is presented on each panel to facilitate comparison.

In agreement with our previous observations (Sluchanko et al, 2017a; Sluchanko et al, 2017c), *Syn*FRP (apparent *M*_W_ – 29.9 kDa) showed only weak interaction with OCP^WT^ (apparent *M*_W_ – 32.0 kDa) in its dark-adapted form (Fig. 4A), but tight interaction occurred with ΔNTE (apparent *M*_W_ – 31.0 kDa; heterocomplexes – 50.6 kDa), representing the OCP variant with non-separated domains but with an uncovered FRP binding site (Sluchanko et al, 2017c), and an analogue of the OCP^R^ form, OCP^AA^ (apparent *M*_W_ – 43.1 kDa; heterocomplexes – 53.3 kDa) (Sluchanko et al, 2017a). Unexpectedly, almost the same pattern was observed for *Ana*FRP (apparent *M*_W_ – 31.7 kDa) and *Amax*FRP (apparent *M*_W_ – 29.8 kDa). Indeed, these proteins showed weak interaction with OCP^WT^ but readily formed complexes with ΔNTE and OCP^AA^. Whereas the binding preferences of *Ana*FRP towards the three OCP forms were almost indistinguishable from those of *Syn*FRP (apparent *M*_W_ of the heterocomplexes with ΔNTE and OCP^AA^ were 50.5 and 54.9 kDa, respectively), pronounced differences were observed for *Amax*FRP. This FRP was clearly able to form complexes with ΔNTE (apparent *M*_W_ of the heterocomplexes – 42.0 kDa) and OCP^AA^ (apparent *M*_W_ of the heterocomplexes – 46.4 kDa), but with much lower apparent *M*_W_ than in the *Syn*FRP and *Ana*FRP cases.

To analyze this unexpected difference in masses of the ΔNTE complexes with *Amax*FRP versus *Ana*FRP (or *Syn*FRP) more accurately, we performed SEC-MALLS experiments by loading two pre-incubated mixtures of ΔNTE with a different molar excess of FRP (Fig. 5). In agreement with Fig. 4F, the ΔNTE+ *Amax*FRP profile contained a peak of the complex and also a small shoulder presumably corresponding to the FRP excess. Supporting the value determined from column calibration (42 kDa), the *M*_W_ distribution across the main peak revealed the mean value of 46 kDa exactly coinciding with the equimolar protein ratio (calculated monomer *M*_W_ are 12.6 kDa for *Amax*FRP and 33.4 kDa for ΔNTE), in line with the previous *in vitro* observations suggesting 1:1 apparent stoichiometry for various OCP-FRP complexes (Moldenhauer et al, 2017b; Sluchanko et al, 2017a; Sluchanko et al, 2017c). When a 3.5-fold excess of *Syn*FRP was mixed with ΔNTE, we observed two peaks with the mean *M*_W_ of 59 and 28 kDa, corresponding to the heterocomplex and the excessive FRP. Surprisingly, the amplitude of the remaining FRP peak was consistent with the notion that more than one FRP equivalent moved to the peak of the heterocomplex, in accord with its *M*_W_ = 59 kDa, implying 1:2 apparent OCP:FRP stoichiometry (Fig. 5). The average *R*_H_ values determined from the light scattering for the complexes with 1:2 and 1:1 apparent stoichiometries were also significantly different (3.82 and 3.32 nm, respectively). These completely unexpected results indicate that, depending on conditions, FRP can form not only 1:1 but also 1:2 complexes with ΔNTE. In the light of this finding, the intermediary *M*_W_ values of the ΔNTE complexes with *Syn*FRP (50.6 kDa) and *Ana*FRP (50.5 kDa) observed in Fig. 4D,E most likely reflect a mixture of 1:1 and 1:2 complexes and, therefore, may indicate that the connection between FRP monomers weakens in the heterocomplexes, as would be characteristic for the transitory state between FRP dimer binding to OCP and ultimate formation of the 1:1 complex between OCP and FRP. By analogy, the smaller size of the OCP^AA^/ *Amax*FRP complexes (Fig. 4I) may indicate the same difference in stoichiometry as seen in the case of ΔNTE. This important novel information specifies the mechanism of the FRP interaction with OCP and suggests that the dimer interface in FRP may not be immediately involved in OCP binding, in contrast to our earlier hypothesis (Sluchanko et al, 2017a).

We conclude that despite clear differences in the hydrodynamic behavior and stoichiometry of the OCP complexes with *Amax*FRP compared to that with *Syn*FRP or *Ana*FRP, FRPs with substantially different amino acid composition are able to specifically interact with various forms of an OCP from another species, which is unexpected and demonstrated here for the first time.

To get more insight into potential differences in the OCP binding mechanism between the analyzed FRP species, we compared their ability to interact with the individual domains of *Synechocystis* OCP. In line with our previous observations, *Syn*FRP was unable to bind to the OCP-NTD (also termed RCP) in either its apo- (Fig. 6A; apparent *M*_W_ – 21.7 kDa) or holoform (Fig. S3), but showed interaction with the carotenoprotein COCP (apparent *M*_W_ – 43.8 kDa; heterocomplexes – 51.7 kDa) corresponding to the dimer of two CTDs of *Synechocystis* OCP containing a single carotenoid (Moldenhauer et al, 2017a), implying that the main FRP binding site should be located on the OCP-CTD. This interaction with COCP is independent of the presence of carotenoid (Moldenhauer et al, 2017a). A similar preference towards COCP was demonstrated by *Ana*FRP, which also did not interact with the RCP apoprotein (Fig. 6B). Neither did *Amax*FRP (Fig. 6C); however, in its case we could barely detect interaction even with COCP (Fig. 6F), in contrast to *Syn*FRP (Fig. 6D) and *Ana*FRP (Fig. 6E). Therefore, *Amax*FRP, which under the conditions used readily interacts with full-length OCP variants by forming exclusively 1:1 complexes (Fig. 4F, I), is virtually incapable of binding individual CTDs in the form of the COCP dimer. These observations may mean that the FRP-binding site on the OCP-CTD is just one part of the (multisite) FRP-binding region in OCP, since all tested FRP species including *Amax*FRP showed interactions with OCP forms containing two domains (either separated or not). This observation is consistent with the previously postulated hypothesis that FRP works as a scaffold bringing the OCP domains together (Lu et al, 2017; Sluchanko et al, 2017a; Sluchanko et al, 2017c). Different FRP binding modes with OCP seems also probable, especially given the recently proposed hypothesis that FRP has two activities in relation to OCP, i.e., it accelerates the OCP^R^→OCP^O^ transition and it detaches OCP from PBs (Thurotte et al, 2017). Taking into account the co-occurrence of a full-length OCP and individual CTD homologues in some cyanobacteria, the possibility of interaction between FRPs and CTDHs warrants separate detailed investigation.

### Functional interaction of the FRP homologues with Synechocystis OCP

As it was noted, FRP may have two distinct roles: (1) it can increase the rate of the OCP^R^→OCP^O^ transition and (2) it can detach OCP from PBs (Thurotte et al, 2017). In order to compare these functional properties of different FRPs, we tested both functions *in vitro*. It should be noted that spectroscopic monitoring of interactions between OCP, FRP and PBs captures a mixture of multiple simultaneously occurring processes, including diffusion- (and concentration)- dependent binding of FRP to free OCP, binding of FRP to the OCP-PBs complexes, spontaneous or FRP-induced OCP^R^→OCP^O^ conversion, and spontaneous or FRP-induced detachment of OCP from PBs (Shirshin et al, 2017). Considering the complexity of these reactions, we sought for experimental settings to isolate specific stages. As reported previously, the accumulation of the active quenching OCP form can be represented by the following set of transitions: OCP^O^(orange compact, inactive) → OCP^RI^(red compact, inactive) → OCP^R^(red, separated domains, functionally active), with asynchronous changes in the carotenoid and protein components (Maksimov et al, 2017c). Taking this into account, by using ΔNTE we analyzed the effect of FRP within the preformed OCP-FRP complexes on the lifetime of the red state (OCP^R^) with separated domains. Then, using the OCP^AA^ double mutant, which is constantly active in the dark and cannot be inactivated by phosphate (Maksimov et al, 2017c) (Fig. S4), we studied the rates of the FRP-induced detachment of OCP from PBs. Finally, we tested if ΔNTE in the red state (equivalent to OCP^R^) can induce PBs fluorescence quenching upon continuous illumination of the sample by actinic light (AL) in the presence of various FRPs.

Fig. 7A shows that all studied FRPs significantly decrease the lifetime of the red active state of ∆NTE comparing to the case when FRP is absent (~ 3300 ms; see Table S3). Surprisingly, *Syn*FRP did not show the best efficiency of accelerating the decay of the red state of its cognate OCP. Rather, *Ana*FRP accelerated the decay of OCP^RI^ almost two times (~ (50 μs)^-1^) compared to the values in the presence of *Syn*FRP or *Amax*FRP (~ (90–100 μs)^−1^). While this indirectly indicates that the ΔNTE/ *Ana*FRP complex provides a strong interaction and the best environment for the restoration of H-bonds between the carotenoid and Tyr-201/Trp-288, the faster decay of the OCP^RI^ intermediate coincides with a reduced quantum yield for full photoconversion into the OCP^R^ state (~ 84 %) as indicated by the lower intermediate plateau between 1 and 10 ms. Nevertheless, *Ana*FRP was also characterized by the slowest OCP^R^→OCP^O^ back conversion compared to *Syn*FRP indicating a compromised ability to reverse the domain separation. These observations suggest that the ability of FRP to serve as a scaffold for the correct NTD-CTD alignment represents a property which is separate from the stabilizing interactions required for re-establishing the proper chromophore-protein interactions. Considering the observed lifetimes of OCP^R^ and assuming OCP^RI^→OCP^O^ as an elementary act, one can estimate that ΔNTE/ *Ana*FRP needs about 3300 attempts to connect the domains, which is much higher comparing to complexes with *Syn*FRP (~ 1550 attempts) and especially *Amax*FRP (~ 990 attempts). As long as the absence of NTE ensures binding of FRPs to the main site in the CTD, such estimations may indicate that different FRPs have different efficiency of interactions with the secondary FRP binding site(s) tentatively located on the NTD, in the framework of the proposed ‘domain-bridging’ FRP activity (Lu et al, 2017; Sluchanko et al, 2017a; Sluchanko et al, 2017c).

**Fig. 7.**
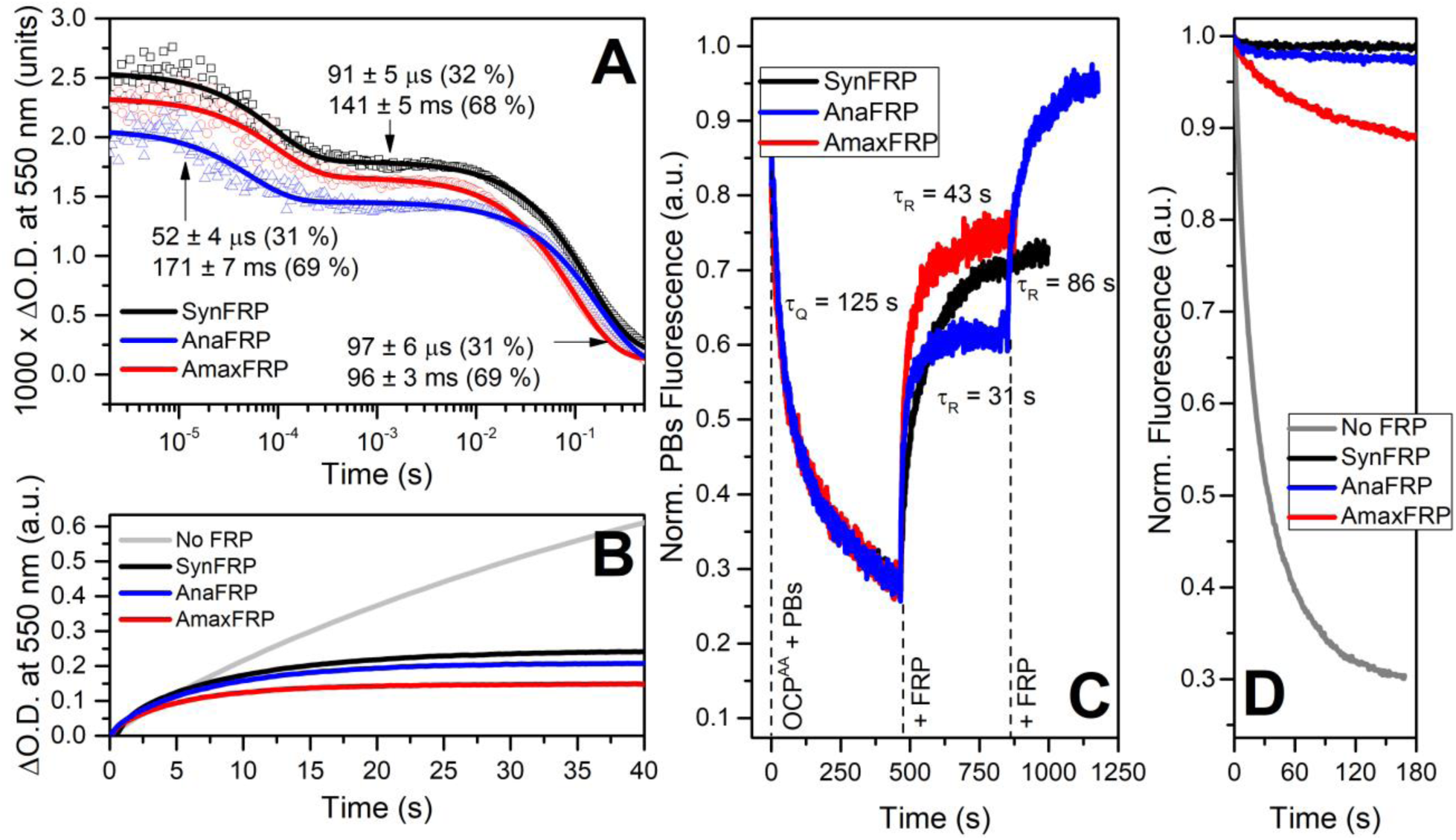
Influence of different FRP species on the OCP photoactivity and the ability to induce PBs fluorescence quenching *in vitro*. (**A**) – flash-induced transitions of ∆NTE measured as changes in optical density at 550 nm in the presence of FRP (2.4 FRP per 1 OCP). Experiment was conducted at 35 °C for increasing the rates of transitions. (**B**) – accumulation of the red form of ∆NTE under continuous illumination of the samples by a blue 200 mW LED in the absence and presence of various FRPs. The experiments were conducted at 5 °C in order to reduce the rates of OCP^R^→OCP^O^ transitions and to reveal effects of different FRPs (see Table S3 and text for more details). (**C**) – time-courses of PBs fluorescence quenching induced by OCP^AA^ in the dark followed by fluorescence recovery upon addition of FRPs. Experiments were conducted at 25 °C and constant stirring. (**D**) – time-courses of PBs fluorescence quenching induced by photoactivation of ΔNTE by 200 mW blue LED in the absence (gray line) and in the presence of FRP (3.6 FRP per 1 ΔNTE). Experiments were conducted at 25 °C and constant stirring.

Fig. 7B shows that, under continuous illumination by AL, OCP^R^ gradually accumulates in the absence of FRPs, whereas their presence significantly reduces the amplitude of the changes and accelerates the onset of the equilibrium state between the processes of OCP^R^ formation and decay (Table S3). An overall reduction of the OCP^R^ concentration in the presence of FRPs could be explained by an increase of OCP^R^→OCP^O^ rate (according to the model reported in (Maksimov et al, 2015)), which was indeed observed in experiments with *Amax*FRP, for which the OCP^R^→OCP^O^ rate is the highest, while corresponding values for *Ana*FRP and *Syn*FRP are comparable (Fig. 7A, Table S3). We may speculate that formation of the red form of ΔNTE in complex with FRP requires interruption of OCP-FRP binding at the secondary site(s) and is thus determined by peculiarity of protein-protein interfaces between OCP and different types of FRPs.

Further, using the constantly active OCP^AA^ mutant we were able to estimate OCP–PBs detachment rates in the presence of FRPs (Fig. 7C). Surprisingly, all studied FRP species induced fluorescence recovery of *Synechocystis* PBs quenched by OCP^AA^; the fastest recovery was observed in the *Ana*FRP case, which, as we suppose, is related to the abovementioned stability of the OCP–FRP complex formation. Unfortunately, at present we do not know if the recovery of PBs fluorescence occurs due to the active detachment of the quencher by FRP or due to the spontaneous breakdown of the dynamically formed OCP^AA^–PBs complexes accompanied by formation of the OCP^AA^–FRP complexes resulting in OCP^AA^ scavenging by FRP, which may prevent further interactions of OCP^AA^ with PBs. It should also be noted that due to the absence of the H-bond donors to the ketocarotenoid in the CTD of OCP^AA^, even the proper positioning of OCP domains assisted by FRP cannot cause formation of the orange form, thus the FRP-induced detachment of OCP from PBs occurs regardless of the spectral characteristics of the OCP state (red *vs.* orange), supporting the existence of several independent functional activities of FRP. If FRP is present in excess, the initial level of PBs fluorescence could be reached (as also indicated by the FRP dose-response shown exemplarily for *Ana*FRP in Fig. 7C), indicating that all constantly active OCP^AA^ molecules are scavenged by FRP.

Surprisingly, not all FRPs were able to completely prevent PBs fluorescence quenching by the red active form of ΔNTE (Fig. 7D) after formation of the active ΔNTE form by AL. This phenomenon is clearly visible in the case of *Amax*FRP, while photoinduced PBs fluorescence quenching with other FRPs is almost negligible. Such a behavior can be explained by several possibilities: (i) – binding of the active form of ΔNTE to PBs is more efficient comparing to OCP^AA^, and (ii) – binding of *Amax*FRP to ΔNTE in a distinctly different stoichiometry compared to other studied FRPs (Fig. 5) may not fully prevent the ability of ΔNTE to quench PBs fluorescence upon photoactivation, in other words, in complex, *Amax*FRP may not fully block the exposure of the interface in the OCP-NTD responsible for interactions with PBs.

## Discussion

Under high light, OCP is reversibly photoconverted to the active but metastable OCPR form, which is considered the main target of FRP binding and action (either in free or PBs-bound OCP state). Upon binding to OCP^R^ with separated domains, FRP accelerates its back conversion to OCP^O^, dramatically decreasing the OCP^R^ lifetime. This makes potentially informative structural studies very challenging and, therefore, the whole process of the FRP-regulated OCP functioning on a molecular level so poorly understood. In this respect, detailed investigation of more kinetically stable intermediates of the OCP photocycle, OCP mutants and individual domains, in complex with FRP seems much more promising. By now, the strongest FRP binders not requiring photoactivation are the purple mutant OCP forms (Maksimov et al, 2017c; Sluchanko et al, 2017a) having key Trp/Tyr residues participating in the H-bonding to carotenoid mutated (apparent *K*_D_ ~2-3 μM, i.e. at least 10 times stronger than for OCP^O^ (Sluchanko et al, 2017a)) and the ΔNTE form with non-separated domains (apparent *K*_D_ <1 μM (Sluchanko et al, 2017c)); the binding is also observed with COCP composed of CTD dimers, but not with RCP or RCP^apo^ (=NTD) (Moldenhauer et al, 2017a; Sluchanko et al, 2017b; Sutter et al, 2013), implying the presence of the main FRP binding site on the OCP-CTD. It was also hypothesized that FRP can act as a scaffold bridging the two OCP domains together (Lu et al, 2017; Sluchanko et al, 2017a) and can have an extended, multisite binding region on OCP (Yang et al, 2015), which is supported by observation of intermediary NTD-FRP-CTD complexes by native mass-spectrometry (Lu et al, 2017) and compaction of the OCP forms with separated domains upon FRP binding (Moldenhauer et al, 2017b; Sluchanko et al, 2017a). Phe-299 (Thurotte et al, 2017) and other hydrophobic residues on the OCP-CTD, covered by NTE in OCP^O^ (Sluchanko et al, 2017c), were proposed to be important for FRP recruitment and action on OCP. Intriguingly, FRP binding can be accompanied by its monomerization, which earlier led to a suggestion about the role of the subunit interface in the interaction process (Sluchanko et al, 2017a; Sluchanko et al, 2017c). Conformational changes involving the unfolding of the head FRP domain were hypothesized to play a critical role in the FRP mechanism (Lu et al, 2017), whereby FRP can serve as a lever arm or jack bringing the domains close to each other. The proposed conformational changes seem very reasonable and may somehow destabilize the subunit interface within the OCP-bound FRP, explaining its monomerization. Nevertheless, the whole series of interconversions, as well as stoichiometries and structures of intermediary complexes formed during the OCP photocycle in the presence of FRP are not clearly elucidated. Moreover, the main mechanistic conclusions have been drawn from studies on *Synechocystis* proteins so far, leaving the question about universality and conservativity of the FRP mechanism among different cyanobacteria dramatically underexplored.

Phylogenetic analysis shows that multiple FRP-like sequences are much less identical than their OCP counterparts. Besides the majority of species containing both OCP and FRP, there is a significant number of cyanobacteria that have only OCP (Bao et al, 2017) without, or along with different number of homologues of its NTD and CTD (Melnicki et al, 2016). Recently, the existence of unusual inducible OCP variants capable of spontaneous relaxation without requiring FRP (termed OCP2, opposite to the more classical OCP1) has been demonstrated and it was hypothesized that OCP2 variants expand adaptational capabilities of the corresponding cyanobacteria (Bao et al, 2017). Surprisingly, there are four cyanobacterium species which have FRP genes, while OCP, HCP and CTDH genes are absent (Bao et al, 2017), implicating that FRP homologues may have roles beyond those associated with OCP.

In the framework of the classic OCP1 system requiring FRP, we selected and characterized two FRP homologues from *A. variabilis* and *A. maxima* having very limited sequence identity with *Syn*FRP. Interestingly, the two analyzed FRPs belong to the OCP/FRP containing group of cyanobacteria ( *Syn*FRP, *Amax*FRP), whereas the third ( *Ana*FRP) belongs to a cyanobacterium having, along with one OCP gene and one FRP gene, also a set of NTD homologues and one CTD homologue (Boulay et al, 2008; Lopez-Igual et al, 2016; Melnicki et al, 2016).

Structural analysis of these previously uncharacterized low-homology proteins by using state-of-the-art techniques reveal a highly similar dimeric conformation in solution (Figs 2 and 3), with the possibility of an angular shift between the subunits that is also to some extent observed in crystals of FRP dimers from *Synechocystis* (PDB 4JDX) and *Tolypothrix* (PDB 5TZ0) (Sluchanko et al, 2017b). Such a sliding of FRP monomers relative to each other suggests that FRP dimers are not rigid entities and it may be relevant for the conformational changes in the OCP-bound FRP and its monomerization whose cause-and-effect relation is not yet clear.

Completely unexpectedly, FRP homologues preserved the preferences of *Syn*FRP towards the studied OCP forms from *Synechocystis* (Fig. 4 and 6, Fig. S3), but the stoichiometries observed in the case of *Amax*FRP heterocomplexes compared to those of other two FRPs were markedly different. Under various conditions used, this FRP formed almost exclusively 1:1 complexes with ΔNTE (and presumably, OCP^AA^), whereas *Syn*FRP and *Ana*FRP could also form 2:1 complexes. This may tentatively indicate that these complexes reflect different intermediary states having distinct stabilities if formed by different FRPs. Intriguingly, only *Amax*FRP was not able to form complexes with COCP, which potentially has two available FRP binding sites per CTD dimer. One explanation may be that, in order to tightly bind to OCP, this particular FRP may require a more expanded binding interface than can be provided by the CTD alone, i.e., requires secondary contacts (in the interdomain linker or NTD) that would be in line with the ‘domain-bridging’ activity of FRP. The remarkable difference in *Amax*FRP binding to the ΔNTE and OCP^AA^ forms of OCP makes this heterologous FRP very interesting and promising OCP partner in structural studies in the future.

The similarity of the structures prompted us to map the surface of a FRP dimer according to the conservativity of various FRP sequences (Fig. 8). In agreement with the data of Sutter et al. (Sutter et al, 2013), the main conserved surface is found in the dimerization region, however, the two other highly conserved sites are located in head domains of FRP, whereas the convex surface is more variable (Fig. 8). It is reasonable to suggest that these immutable, evolutionary cold spots can be responsible for the FRP functioning and its universality. The potential role of the dimerization region in binding to OCP has already been discussed and supported by mutational studies (Sutter et al, 2013). The importance of the conserved region located in the head domains of FRP is less understood; however, the replacement of a highly conserved Phe-76 and Lys-102 from this region ( *Synechocystis* numbering) severely affects the FRP-OCP interaction (Lu et al, 2017), commensurate with the hypothesis about the role of the conserved region in head domain.

**Fig. 8.**
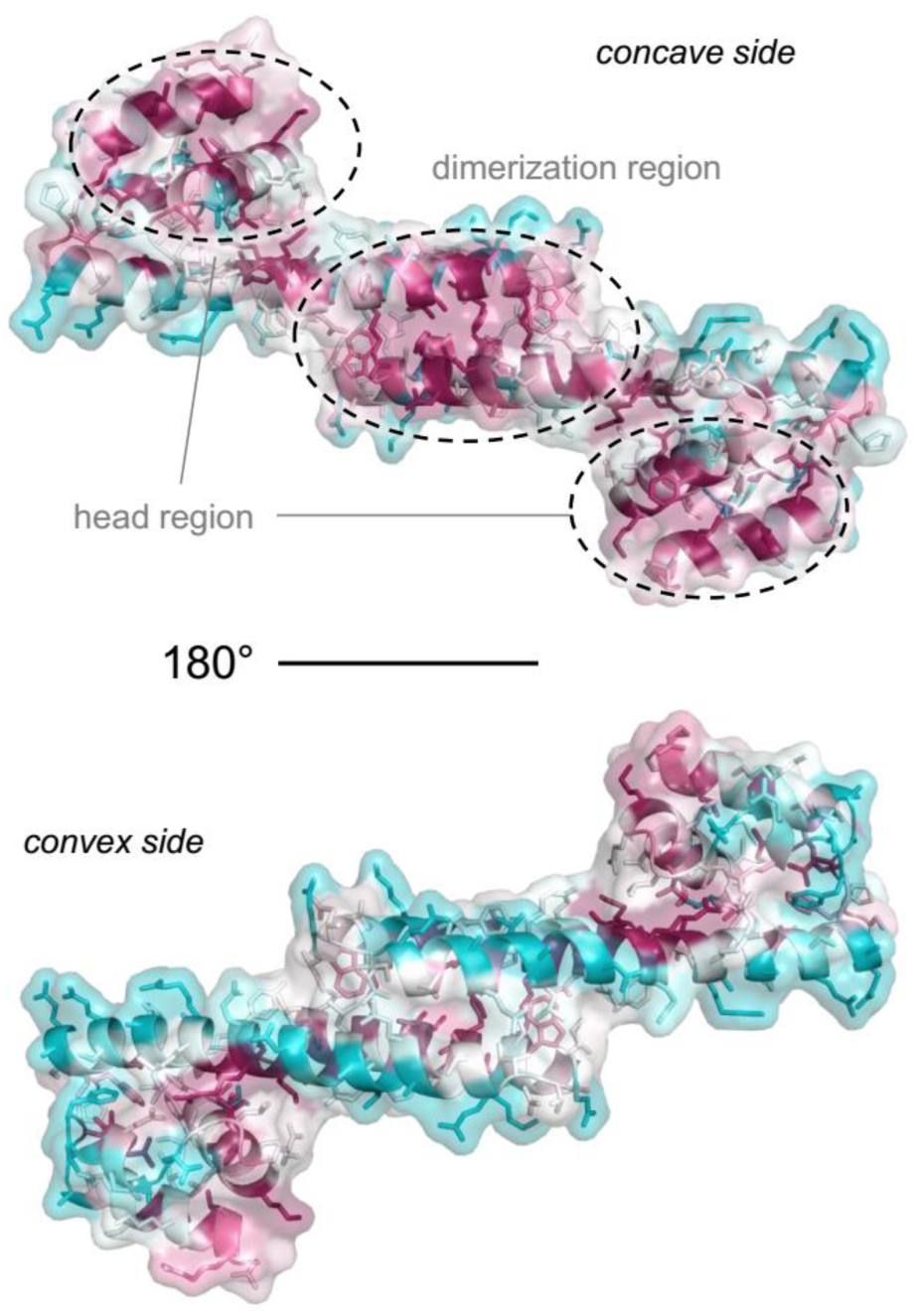
Conservativity analysis performed using all known FRP homologues using *CONSURF* (Ashkenazy et al, 2016) showing the most conserved regions in the dimerization and head domains of the FRP structure (represented in two projections). The gradient from the most conserved (purple) to the most variable (cyan) is used.

Functional tests showed that the selected low-homology FRPs do perform on *Synechocystis* OCP and influence various aspects of its photoprotecting function, confirming a certain level of universality of the FRP mechanism. Indeed, all FRPs were able to accelerate the OCP^R^→OCP^O^ back conversion, to reduce accumulation of the OCP^R^ form under AL (and speed up the achievement of the equilibrium state), to recover PBs fluorescence by detaching the PBs-bound OCP, and to prevent OCP-induced quenching of PBs (Fig. 7). At the same time, the recently accumulated knowledge and the ability to accurately assess the effects of FRPs on different intermediates of the OCP photocycle (Maksimov et al, 2017c) and on specific OCP forms (OCP^R^ analogues (Maksimov et al, 2017c; Sluchanko et al, 2017a) and ΔNTE (Maksimov et al, 2017c; Sluchanko et al, 2017c)) helped us to reveal important mechanistic differences. The most surprising observations are related to *Amax*FRP, which was capable of the most efficient conversion of OCP^R^ into OCP^O^, and detached OCP from PBs comparing to *Syn*FRP at least twice faster, but still was not able to completely prevent PBs fluorescence quenching under AL. These facts strongly suggest that PBs-OCP and OCP-FRP complexes should be considered as a metastable structure. Further investigations of this phenomenon are of particular interest. Together with interaction studies (Figs 4-6), our functional analyses (Fig. 7) support the idea that there is more than one FRP-binding interface on OCP (one is definitely located in the CTD and the second one(s), presumably, in the NTD) and suggest that heterologous FRPs may display different affinity towards the main and the secondary FRP binding site, representing highly useful tools to probe the FRP-mediated mechanism.

Thus, the present study makes the first step to understand the universality and conservativity of the FRP mechanism in the OCP-mediated photoprotection system of cyanobacteria, and future research using other FRP and OCP homologues should expand the findings reported here. We expect that utilization of different FRP homologues may also shed new light on the mechanistic aspects of FRP functioning and will be helpful for structural studies in the future.

## Materials and methods

### Protein cloning, expression and purification

Cloning, expression and purification of the His_6_-tagged *Synechocystis* RCP and FRP were described previously (Moldenhauer et al, 2017a; Sluchanko et al, 2017a). The cDNA sequence for the ‘constantly quenching’ OCP^Y201A/W288A^ mutant protein ((Maksimov et al, 2017c); termed OCP^AA^ in this study) was generated using the QuikChange site-directed mutagenesis kit and cloned into the pQE81L plasmid (amplicillin resistance) by *BamHI*/ *NotI* restriction sites. To permit truncation of flexible N-termini including the His_6_ tag, a cleavage site for the highly specific human rhinovirus 3C protease (recognition amino acid sequence LEVLFQ/GP) was inserted immediately upstream of the endogenous Pro-2 or Pro-13 in the *Synechocystis* OCP sequence, which after 3C protease cleavage produced the constructs OCP_2-317_ (termed OCP^WT^ herein, N-terminal amino acid sequence GP(2)FTIDSARGI…), OCP_13-317_(equivalent to and termed ΔNTE herein, amino acid sequence GP(13)NTLAADVVP…). The 3C cleavage site was also inserted into the plasmid harboring the cDNA of the C-terminal domain of *Synechocystis* OCP, yielding after 3C cleavage the N-terminal amino acid sequence: GPDPATA(165)GKDGKRIAE… (construct corresponding to residues 165-317). For obtaining *Synechocystis* FRP_8-109_, the 3C site was introduced before Pro-9 yielding after cleavage the amino acid sequence GP(9)WSQAETQSA…. cDNA sequences were subcloned into the pRSFDuet-1 plasmid (kanamycin resistance) via *BamHI*/ *NotI* restriction sites. cDNA sequences of *Arthrospira* FRP [Uniprot entry B5W3T4 ( *Arthrospira maxima* CS-328), coincides with Uniprot entries H1W9V5 ( *Arthrospira* sp. PCC 8005) and K1X0E1 ( *Arthrospira platensis* C1)] and *Anabaena* FRP [Uniprot Q3M6D9 ( *Anabaena variabilis* PCC 7937), coincides with Uniprot entry A0A1W5CLT8 ( *Anabaena* sp. 39858)] were obtained by artificial gene synthesis (GeneArt, Regensburg, Germany; sequences available upon request) and subcloned into an appropriately modified pQE81L plasmid (harboring a 3C cleavage site before the start methionine) via *BamHI*/ *NotI* restriction sites. The identity of cDNAs was verified by DNA sequencing (Eurofins MWG Operon, Ebersberg, Germany).

Holoforms of OCP^WT^, ΔNTE, RCP, COCP and OCP^AA^ were expressed in echinenone (ECN) and canthaxanthin (CAN)-producing *E. coli* cells essentially as described before (Maksimov et al, 2016; Maksimov et al, 2017b). All His_6_-tagged proteins were purified by immobilized metal-affinity and size-exclusion chromatography (IMAC and SEC, respectively) to electrophoretic homogeneity and stored at +4 °C in the presence of 3 mM sodium azide. Protein concentrations were determined spectrophotometrically using calculated molar extinction coefficients according to Supplementary Table S2. The obtained holoprotein preparations exhibited visible-to-UV absorption ratios of 1.6–1.8 (in case of COCP – 2.5), indicating high sample purity with respect to the contaminating apoprotein.

After IMAC purification, fractions containing target protein were digested using His_6_-tagged 3C protease during dialysis at 4 °C against 1 L of 20 mM Tris-HCl buffer (pH 7.6) containing 300 mM NaCl, 0.1 mM ethylenediaminetetraacetate (EDTA), 0.1 mM phenylmethylsulfonyl fluoride (PMSF), 1 mM dithiothreitol (DTT). The dialysate was clarified by centrifugation for 20 min at 12,000 g and then subjected to the second IMAC to remove 3C protease. The collected protein fractions were combined and the samples were finally purified by SEC.

Phycobilisomes were obtained from *Synechocystis* cells as described previously (Sluchanko et al, 2017a).

### Analytical SEC

To study concentration dependences of hydrodynamics of proteins and the interaction of FRP homologues with either OCP^WT^, ΔNTE, OCP^AA^, or individual OCP domains (RCP and COCP, respectively) we used analytical size-exclusion chromatography (SEC) on two different Superdex 200 Increase (GE Healthcare) columns: 10/300 or 5/150. The smaller column (5/150) allowed long series of experiments to be done under identical conditions in one day to ensure the best data comparison. Protein samples were pre-incubated for at least 15 min at room temperature and then separated using either column equilibrated with a 20 mM Tris-HCl buffer, pH 7.6, containing 150 mM NaCl, 0.1 mM EDTA, and 3 mM ME and calibrated using bovine serum ablumin (BSA) monomer (66 kDa), BSA dimer (132 kDa), BSA trimer (198 kDa), and α-lactalbumin monomer (15 kDa). Flow rates are specified in each particular case. The elution profiles were followed simultaneously by 280-nm and carotenoid-specific absorbance (wavelengths are specified in the respective figure legends). Typical results obtained in at least three independent experiments are presented.

The absolute masses of the ΔNTE complexes with either *Amax*FRP or *Syn*FRP were analyzed on a Superdex 200 Increase 10/300 column using multiparametric detection. Multi-angle laser light scattering (MALLS) with dynamic light scattering (DLS) data were measured in parallel using a Wyatt Technologies Mini-Dawn TREOS with inbuilt quasi-elastic light scattering (QELS) module coupled to a OptiLab T-Rex refractometer for protein concentration determination (dn/dc was taken as 0.185). The MALLS system was calibrated relative to the scattering from toluene and, in combination with concentration estimates obtained from RI, was used to evaluate the *M*_W_ distribution of species eluting from the SEC column. The molecular weight estimates from MALLS/RI and the *R*_H_ derived from DLS were determined using Wyatt ASTRA7 software.

### Small angle X-ray scattering (SAXS) data collection and processing

SAXS data ( *I*( *s*) vs *s*, where *s* = 4πsin *θ*/λ, 2 *θ* is the scattering angle and λ=1.24 Å) from samples of truncated *Synechocystis* FRP ( *Syn*FRP_8-109_, residues 8–109) or full-length *Arthrospira* FRP ( *Amax*FRP, residues 1–106) were measured at the EMBL P12 beam line (PETRA III, DESY Hamburg, Germany; (Blanchet et al, 2015)) using a batch mode (for *Amax*FRP) or the inline SEC-HPLC system (for *Syn*FRP_8-109_) coupled to the MALLS/DLS/RI detectors described above in a common matched buffer (20 mM Tris-HCl buffer (pH 7.6) containing 150 mM NaCl, 0.1 mM EDTA, 2 mM DTT, and 3 % v/v glycerol; 20 °C). The batch mode SAXS data collected from *Amax*FRP (1 s exposure time, collected as 20 x 50 ms frames) at the sample concentrations 1.2–5.8 mg/ml (91–460 μM per monomer) showed little concentration dependence and the data obtained at the highest concentration (460 μM) were used for further analysis. For SEC-SAXS, 100 μl of *Syn*FRP_8-109_ was loaded at high concentration (460 μM) onto a Superdex 200 Increase 10/300 column (GE Healthcare) and eluted at a flow rate of 0.5 ml/min. The flow was equally divided between the SAXS measurements (3600 x 1 s frames) and the MALLS/DLS/RI detection modules (Graewert et al, 2015) to ensure parallel data collection from equivalent parts of the elution profile. For both the batch- and SEC-SAXS, the data reduction, radial averaging and statistical analysis (e.g., to detect radiation damage, or scaling issues between frames) were performed using the *SASFLOW* pipeline (Franke et al, 2012). Statistically similar SAXS profiles were averaged and the buffer scattering subtracted to produce *I*( *s*) vs *s* scattering profiles for *Amax*FRP and *Syn*FRP_8-109_. The SEC-SAXS data were processed using *CHROMIXS* (Panjkovich & Svergun, 2017). *ATSAS* 2.8 (Franke et al, 2017) was employed for the data analysis and modelling. The program *PRIMUS* (Konarev et al, 2003) was used to perform Guinier analysis from which the radius of gyration, *R* _*g*_, and extrapolated zero-angle scattering, *I*(0), were determined (ln *I*( *s*) versus *s*^2^ that were linear in the *sR* _*g*_ range reported in Table S1). The probable frequency of real-space distances, or *p*( *r*) distributions, were calculated using *GNOM* (Svergun, 1992) that provided additional *R* _*g*_ and *I*(0) estimates and the maximum particle dimension, *D* _*max*_. The Porod volume, subsequent hydrodynamic parameters and concentration-dependent and independent *M*_W_ estimates of *Amax*FRP and *Syn*FRP_8-109_ are presented in Table S1.

### Modelling of the solution conformation of SynFRP and AmaxFRP dimers

The *ab initio* bead modelling of both proteins was done using *GASBOR* (Svergun et al, 2001) while *SASREF* (Petoukhov & Svergun, 2005) was used to rigid-body refine the crystallographic structure of *Syn*FRP_8-109_ (PDB 4JDX) to the SAXS data. The atomistic model of *Amax*FRP monomer (residues 1–106) was built using *iTASSER* (Yang et al, 2015) with default parameters; the top scoring model was then aligned to *Syn*FRP subunits to generate *Amax*FRP dimer. Modelled scattering intensities from either the *SASREF* model of *Syn*FRP_8-109_, the *iTASSER* model of *Amax*FRP or the related *Tolypothrix* FRP homologue (PDB 5TZ0) were calculated using *CRYSOL* (Svergun et al, 1995). All data-model fits, as well as the reciprocal-space fit of *p*( *r*) and pair-wise frame comparisons, were assessed using the reduced *χ*^2^ test and Correlation Map (CorMap) *P*-value, whereby *χ*^2^ of ~1 and a CorMap *P* > 0.05 indicate no systematic discrepancies (Franke et al, 2015). CorMap values are reported in Supplementary Table S1. The final SAXS models were deposited to SASBDB (Valentini et al, 2015) under the accession codes SASDD42 ( *Syn*FRP_8-109_) and SASDD52 ( *Amax*FRP). Structural models were drawn in *PyMOL*.

### Absorption spectroscopy

Steady-state absorption spectra, kinetics and 7-ns 532-nm laser flash-induced transients were recorded as described in (Maksimov et al, 2017c). PBs fluorescence quenching was measured as described in (Sluchanko et al, 2017a). Upon absorption and fluorescence measurements, a blue light-emitting diode (LED) (M455L3, Thorlabs, USA), with a maximum emission at 455 nm was used for the photoconversion of the samples (actinic light for OCP^O^→OCP^R^ photoconversion). Temperature of the sample was stabilized by a Peltier-controlled cuvette holder Qpod 2e (Quantum Northwest, USA) with a magnetic stirrer.

## Acknowledgements

The authors are grateful to Prof. Diana Kirilovsky for critical reading of the manuscript and valuable comments. This study was supported by Russian Scientific Foundation project № 18-44-04002. T.F. and M.M. acknowledge support by the German Federal Ministry of Education and Research (WTZ-RUS grant 01DJ15007) and the German Research Foundation (Cluster of Excellence “Unifying Concepts in Catalysis” and FR 1276/5-1). The work was supported by Russian Foundation for Basic Research (grant no. 18-04-00691 N.N.S) and the Horizon 2020 programme of the European Union, iNEXT (653706; D.I.S). N.N.S. thanks iNEXT for the support of the SAXS-683 experiment in EMBL-Hamburg, DESY (iNEXT proposal PID: 2977).

## Author contributions

NNS – conceived the idea and supervised the study; YBS, EGM, TF, NNS – designed the experiments; YBS, EGM, MM, NNS – performed the experiments; CMJ, NNS – performed SAXS data collection and analysis; EGM, CMJ, DIS, TF, NNS – analyzed data; NNS wrote the paper with an input from YBS, EGM, CMJ, DIS, TF.

## SUPPLEMENTARY INFORMATION

**Table S1.**
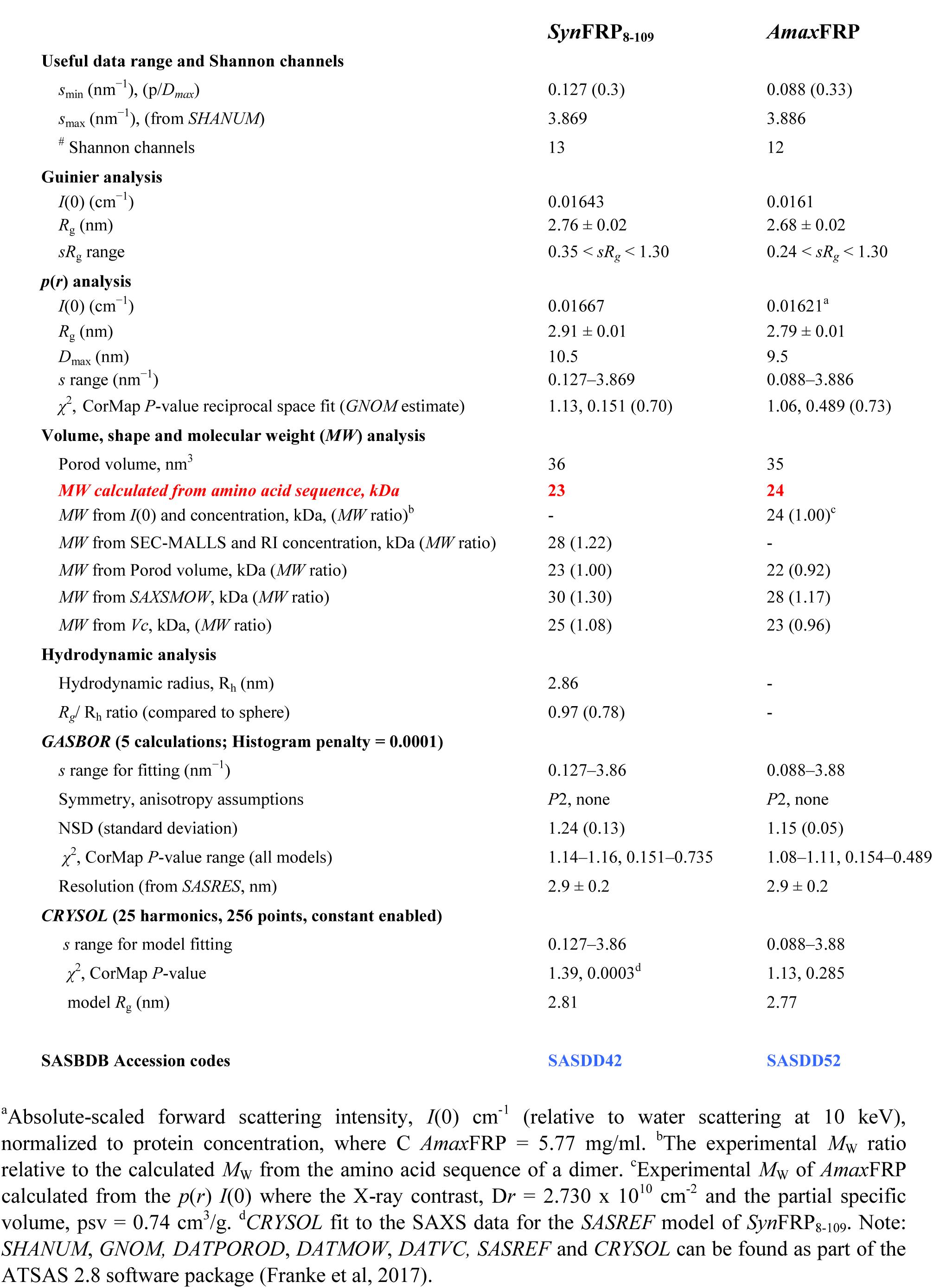
Hydrodynamic parameters determined by SAXS.

**Table S2.** Molar extinction coefficients used in this study for protein concentration determination.

| Abbreviation | Description | $\epsilon_{280}$,<br>$M^{-1}cm^{-1}$ |
| --- | --- | --- |
| *SynFRP* | His-tagged WT (residues 1-109) or untagged (residues 8-109) | 13980 |
| *AnaFRP* | residues 1-108 | 22460 |
| *AmaxFRP* | residues 1-106 | 20970 |
| OCP | residues 2-317 | 34950 |
| $\Delta NTE$ | residues 13-317 | 34950 |
| OCP<sup>AA</sup> | His-tagged; residues 1-317 | 27960 |
| COCP | residues 165-317 | 12490 |
| RCP | His-tagged; residues 1-164 | 22460 |

**Table S3.** Functional interaction of FRP homologues with *Synechocystis* OCP. Lifetimes of OCP^RI^ and OCP^R^ were determined after excitation of the sample by 7 ns laser pulses at 35 °C. Time constants of accumulation of OCP^R^ and corresponding amplitudes of photoconversion were measured at 5 °C under continuous illumination of the samples by AL. PBs quenching experiments were conducted at 25 °C.

| Sample | OCP<sup>RI</sup> →<br>OCP<sup>O</sup>,<br>$\mu s$ | OCP<sup>R</sup> →<br>OCP<sup>O</sup>,<br>ms | Accumulation<br>of OCP<sup>R</sup>, s | Amplitude of<br>photoconversion,<br>% | Photoinduced PBs<br>fluorescence<br>quenching by<br>$\Delta NTE$, % | PBs<br>fluorescence<br>recovery, s |
| --- | --- | --- | --- | --- | --- | --- |
| Control | 90 ± 12 | ~ 3300 | 44.3 | 100 | 100 | - |
| + *SynFRP* | 91 ± 5 | 141 ± 5 | 7.76 | 24.5 | 2.2 ± 1.9 | 86 |
| + *AnaFRP* | 51 ± 4 | 171 ± 7 | 7.03 | 21.0 | 5.3 ± 2.1 | 31 |
| + *AmaxFRP* | 97 ± 6 | 96 ± 3 | 5.31 | 14.8 | 21.0 ± 1.7 | 43 |

**Fig. S1.**
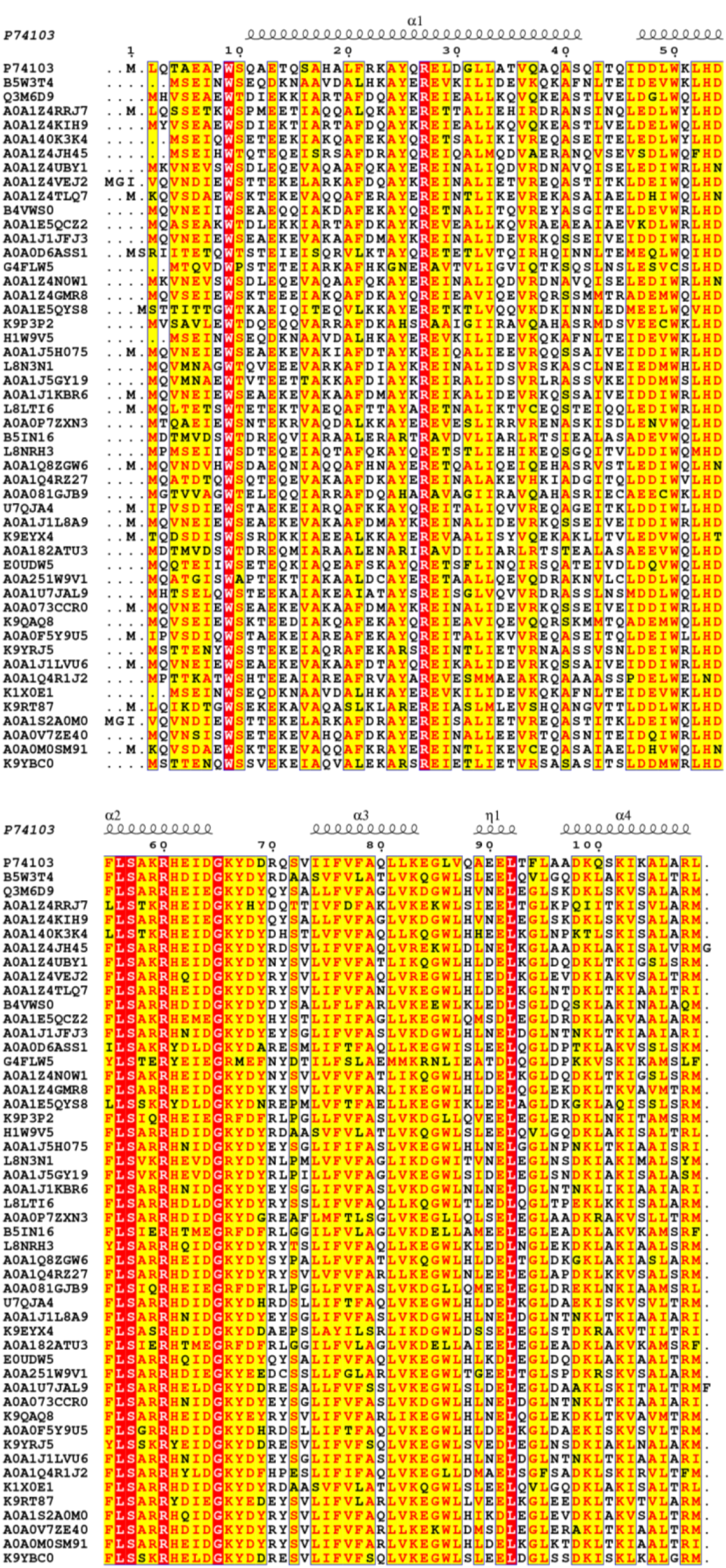
Multiple sequence alignment (MSA) of 50 different FRP-like protein sequences found by a BLAST search using *Synechocystis* FRP (Uniprot P74103) as an entry. The MSA was calculated using M-Coffee (Di Tommaso et al, 2011); representation using the ENDscript server (Robert & Gouet, 2014) shows the assignment of secondary structures (retrieved from the crystal structure of *Synechocystis* FRP) and uses the similarity colouring scheme considering physico-chemical properties of amino-residues. Identical residues are red colored, similar ones – yellow colored. The average consistency of the MSA obtained is 99/100 indicating high robustness of the alignment.

**Fig. S2.**
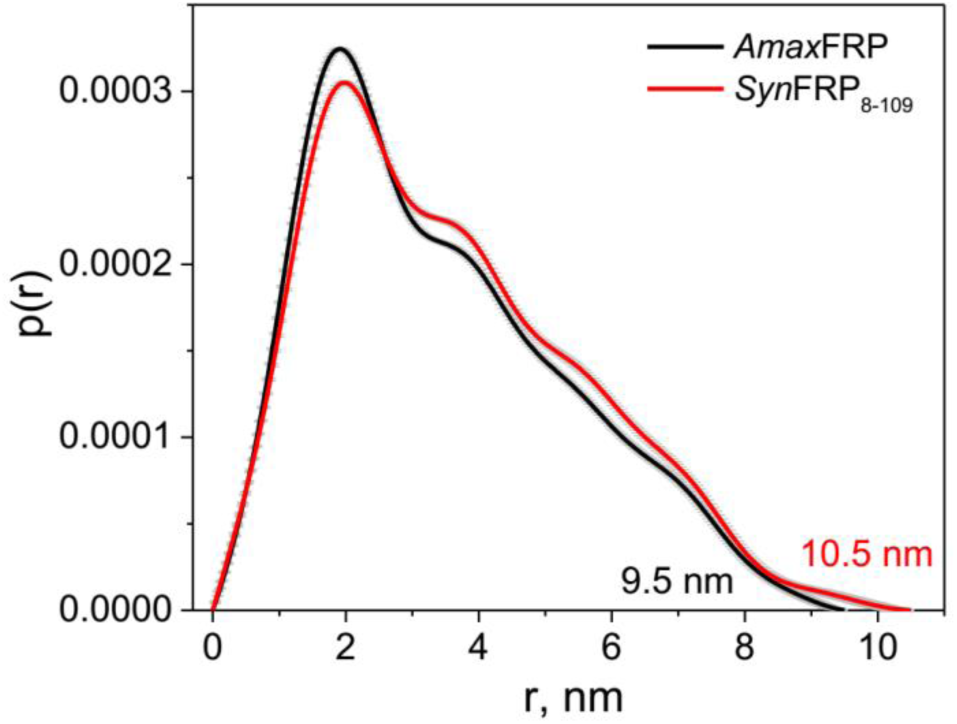
Pairwise distance distribution functions for *Syn*FRP_8-109_ and *Amax*FRP determined using *GNOM* (Svergun, 1992).

**Fig. S3.**
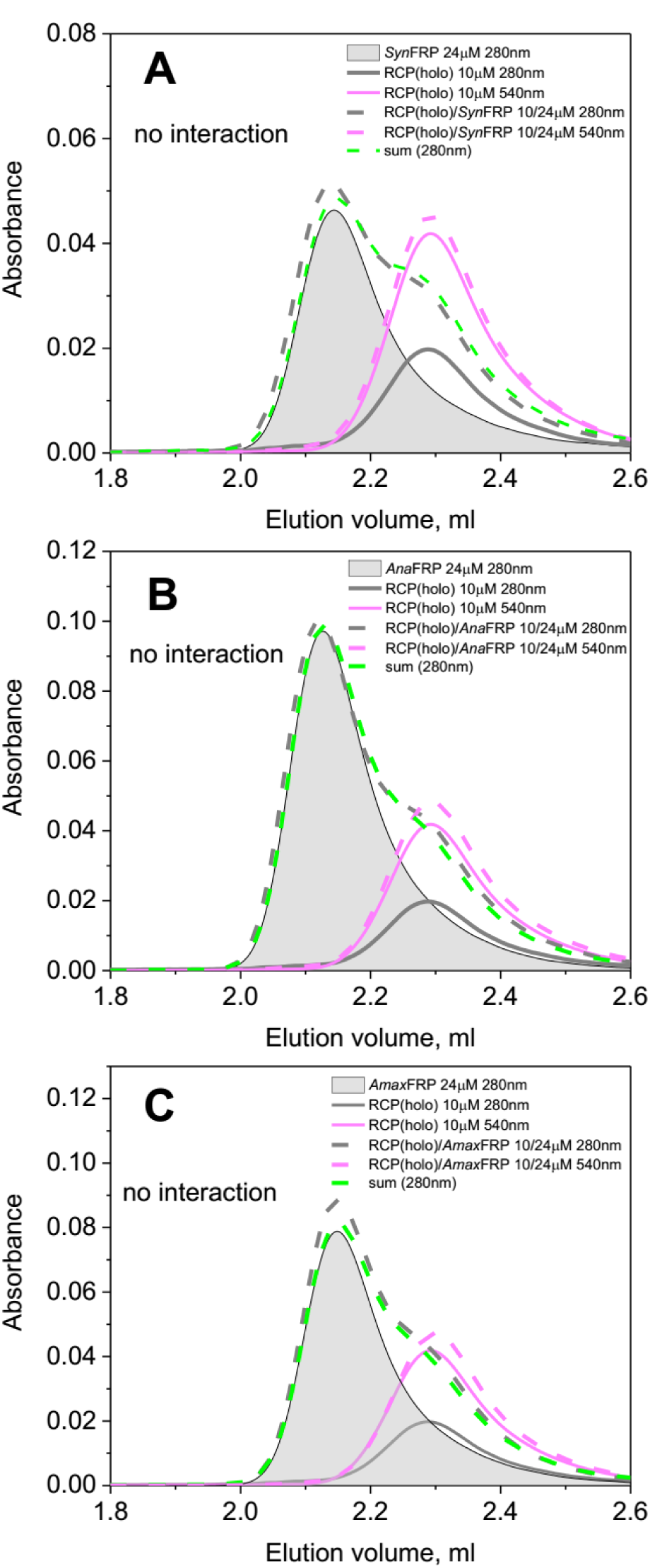
Analysis of the possible interaction between *Syn*FRP (**A**), *Ana*FRP (**B**), or *Amax*FRP (**C**) and holo-RCP using SEC. The samples (40 μl) containing either holo-RCP (10 μM), FRP species (24 μM; semitransparent gray peaks) or their mixtures (dashed magenta and gray lines) at indicated protein concentrations were run on the pre-equilibrated Superdex 200 Increase 5/150 column followed by 280 nm (gray lines) and 540 nm (magenta lines) at a flow rate of 0.45 ml/min. The algebraic sum of the individual 280-nm elution profiles is presented on each panel (light green dashed lines) to facilitate comparison.

**Fig. S4.**
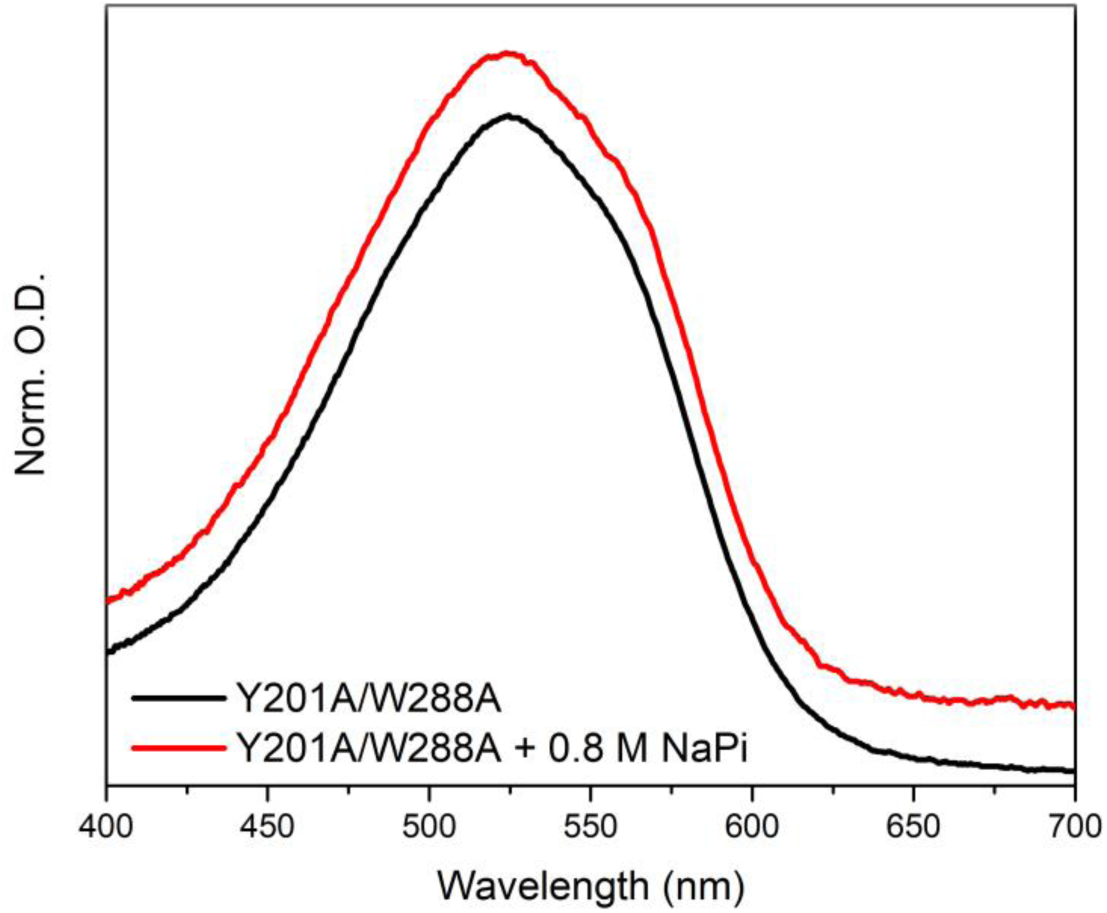
Absorption spectra of the Y201A/W288A mutant of OCP in buffers with low and high concentration of phosphate. For better presentation spectra are shifted along the Y axis. No signatures of the orange form were observed at a high phosphate concentration, indicating that the absence of both residues involved in the formation of H-bonds with the keto-group of carotenoid prevents formation of the basal inactive orange state, in contrast to the single substitution W288A showing substantial “oranging” already in 0.5 M phosphate (Maksimov et al. 2017c). Thus, under experimental conditions suitable for PBs fluorescence measurements the Y201A/W288A mutant is uniquely present in the active form.

**Supplementary text 1.**
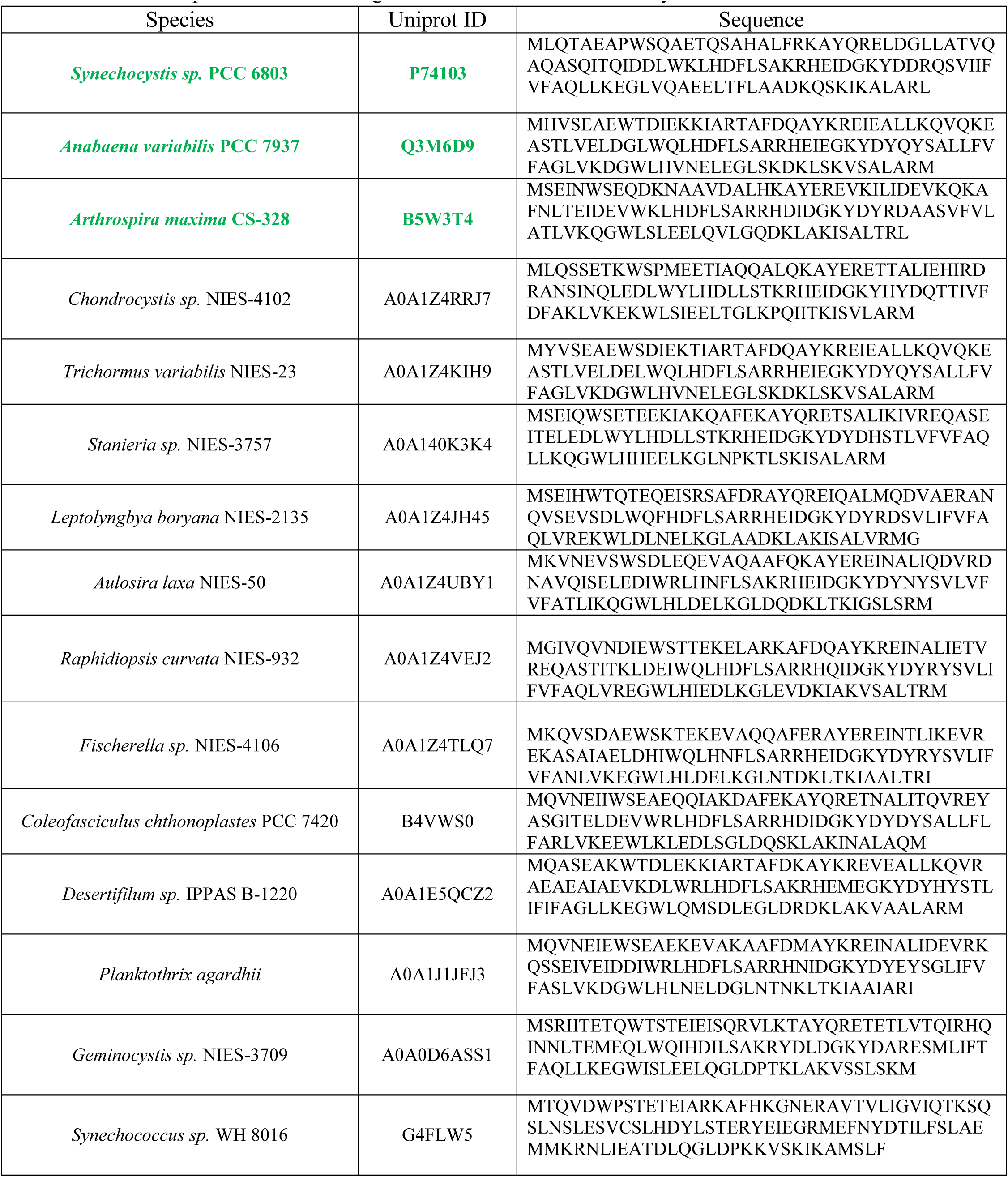
Sequences of 50 FRP-like proteins used to build the MSA (Fig. S1) and phylogenetic tree (Fig. 1A). The sequences were obtained by a BLAST search using default parameters and *Synechocystis* FRP as an entry. Three FRP species marked with green were selected for this study.

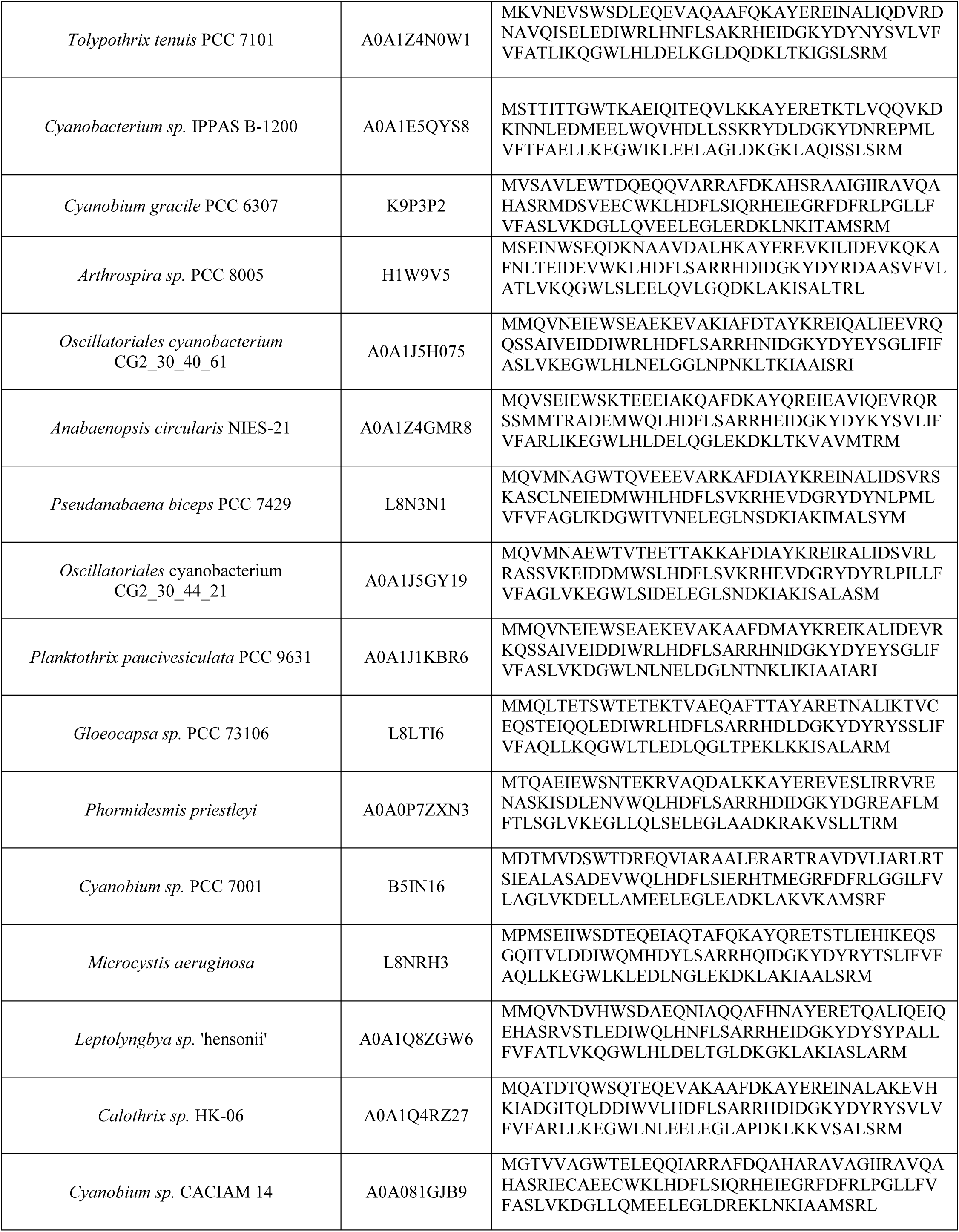

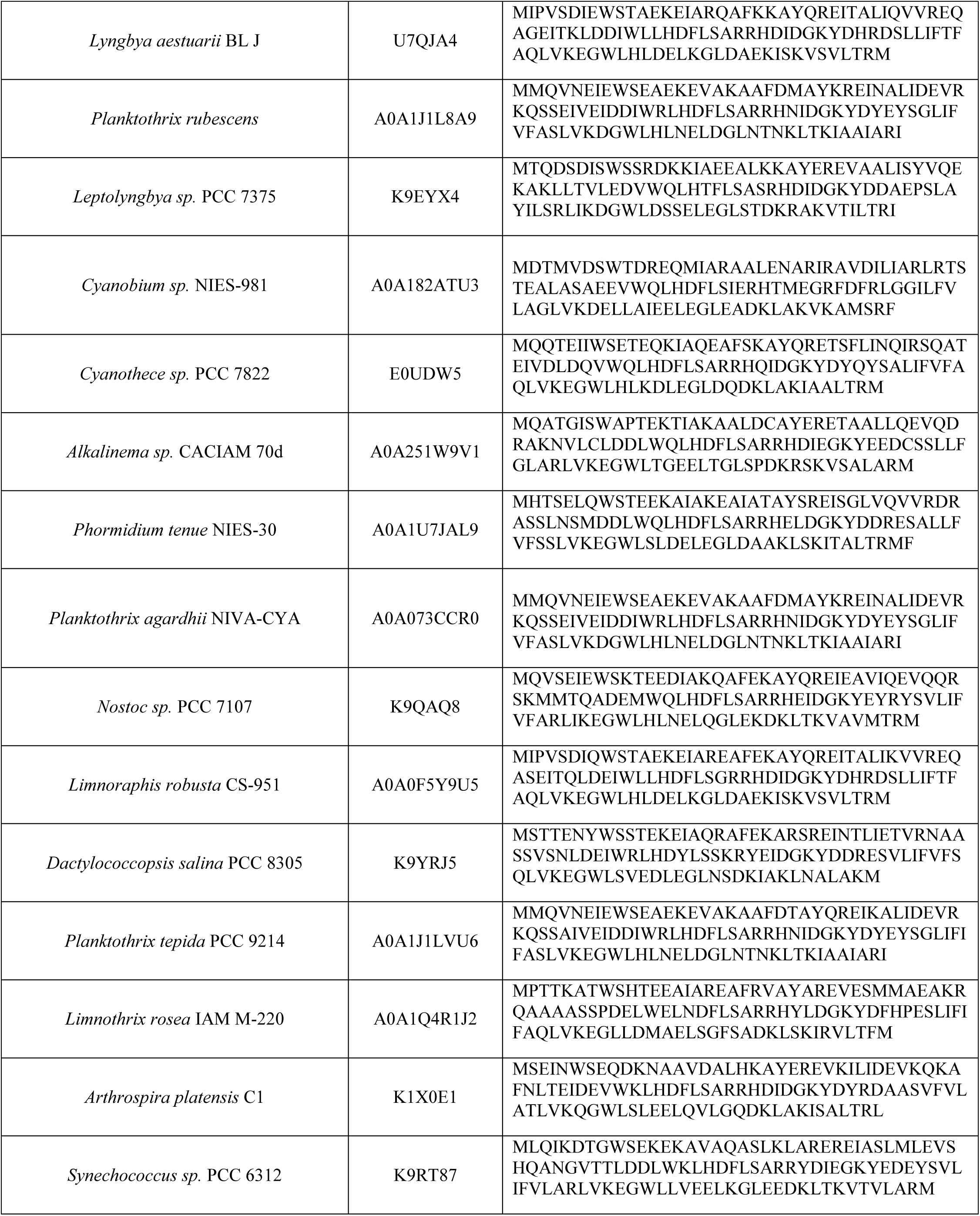

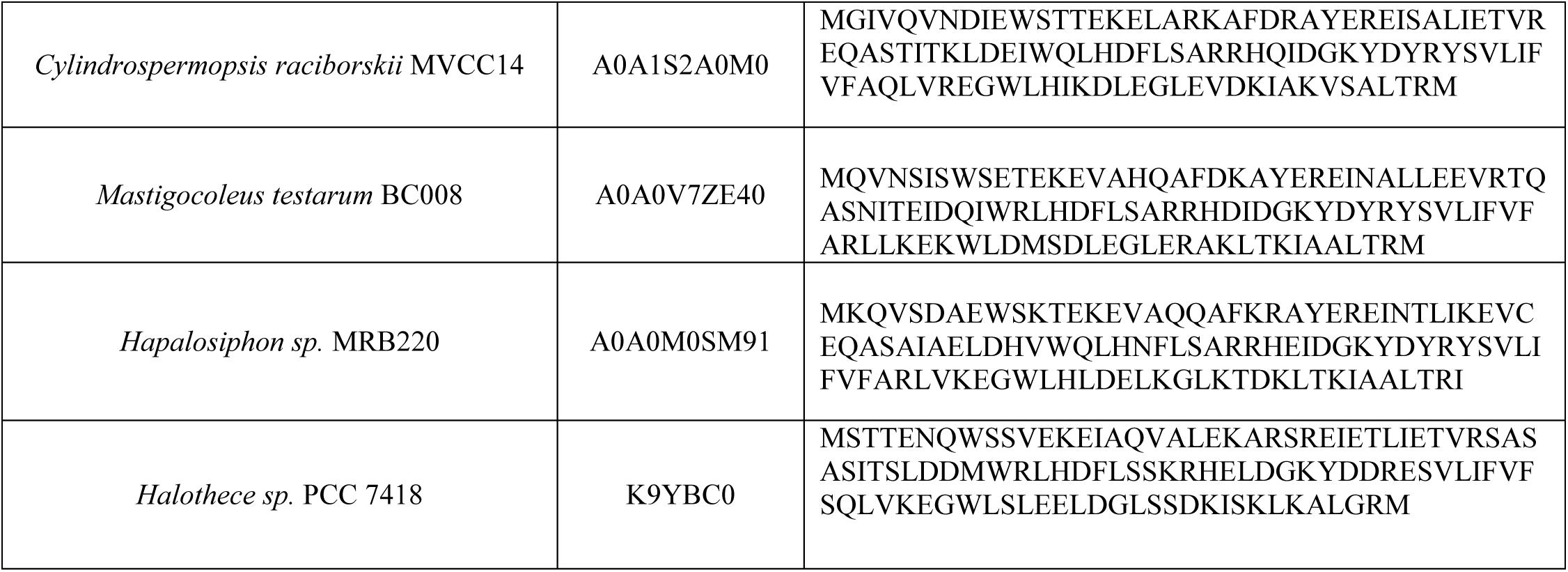

## References

1. Adir N (2005) Elucidation of the molecular structures of components of the phycobilisome: reconstructing a giant. Photosynth Res 85: 15–32

2. Ashkenazy H, Abadi S, Martz E, Chay O, Mayrose I, Pupko T, Ben-Tal N (2016) ConSurf 2016: an improved methodology to estimate and visualize evolutionary conservation in macromolecules. Nucleic Acids Res 44: W344–350

3. Bandara S, Ren Z, Lu L, Zeng X, Shin H, Zhao KH, Yang X (2017) Photoactivation mechanism of a carotenoid-based photoreceptor. Proc Natl Acad Sci U S A 114: 6286–6291

4. Bao H, Melnicki MR, Pawlowski EG, Sutter M, Agostoni M, Lechno-Yossef S, Cai F, Montgomery BL, Kerfeld CA (2017) Additional families of orange carotenoid proteins in the photoprotective system of cyanobacteria. Nature plants 3: 17089

5. Blanchet CE, Spilotros A, Schwemmer F, Graewert MA, Kikhney A, Jeffries CM, Franke D, Mark D, Zengerle R, Cipriani F, Fiedler S, Roessle M, Svergun DI (2015) Versatile sample environments and automation for biological solution X-ray scattering experiments at the P12 beamline (PETRA III, DESY). J Appl Crystallogr 48: 431–443

6. Boulay C, Abasova L, Six C, Vass I, Kirilovsky D (2008) Occurrence and function of the orange carotenoid protein in photoprotective mechanisms in various cyanobacteria. Biochim Biophys Acta 1777: 1344–1354

7. Boulay C, Wilson A, D'Haene S, Kirilovsky D (2010) Identification of a protein required for recovery of full antenna capacity in OCP-related photoprotective mechanism in cyanobacteria. Proc Natl Acad Sci U S A 107: 11620–11625

8. Di Tommaso P, Moretti S, Xenarios I, Orobitg M, Montanyola A, Chang JM, Taly JF, Notredame C (2011) T-Coffee: a web server for the multiple sequence alignment of protein and RNA sequences using structural information and homology extension. Nucleic Acids Res 39: W13–17

9. Felsenstein J (1985) Confidence Limits on Phylogenies: An Approach Using the Bootstrap. Evolution; international journal of organic evolution 39: 783–791

10. Franke D, Jeffries CM, Svergun DI (2015) Correlation Map, a goodness-of-fit test for one-dimensional X-ray scattering spectra. Nat Methods 12: 419–422

11. Franke D, Kikhney AG, Svergun DI (2012) Automated acquisition and analysis of small angle X-ray scattering data. Nuclear Instruments and Methods in Physics Research Section A: Accelerators, Spectrometers, Detectors and Associated Equipment 689: 52–59

12. Franke D, Petoukhov MV, Konarev PV, Panjkovich A, Tuukkanen A, Mertens HDT, Kikhney AG, Hajizadeh NR, Franklin JM, Jeffries CM, Svergun DI (2017) ATSAS 2.8: a comprehensive data analysis suite for small-angle scattering from macromolecular solutions. J Appl Crystallogr 50: 1212–1225

13. Graewert MA, Franke D, Jeffries CM, Blanchet CE, Ruskule D, Kuhle K, Flieger A, Schafer B, Tartsch B, Meijers R, Svergun DI (2015) Automated pipeline for purification, biophysical and x-ray analysis of biomacromolecular solutions. Sci Rep 5: 10734

14. Gupta S, Guttman M, Leverenz RL, Zhumadilova K, Pawlowski EG, Petzold CJ, Lee KK, Ralston CY, Kerfeld CA (2015) Local and global structural drivers for the photoactivation of the orange carotenoid protein. Proc Natl Acad Sci U S A 112: E5567–5574

15. Gwizdala M, Wilson A, Kirilovsky D (2011) In vitro reconstitution of the cyanobacterial photoprotective mechanism mediated by the Orange Carotenoid Protein in Synechocystis PCC 6803. Plant Cell 23: 2631–2643

16. Gwizdala M, Wilson A, Omairi-Nasser A, Kirilovsky D (2013) Characterization of the Synechocystis PCC 6803 Fluorescence Recovery Protein involved in photoprotection. Biochim Biophys Acta 1827: 348–354

17. Holt T, Krogmann DW (1981) A carotenoid-protein from cyanobacteria. Biochimica et Biophysica Acta (BBA) - Bioenergetics 637: 408–414

18. Jones DT, Taylor WR, Thornton JM (1992) The rapid generation of mutation data matrices from protein sequences. Computer applications in the biosciences : CABIOS 8: 275–282

19. Karapetyan NV (2007) Non-photochemical quenching of fluorescence in cyanobacteria. Biochemistry (Mosc) 72: 1127–1135

20. Kerfeld CA, Sawaya MR, Brahmandam V, Cascio D, Ho KK, Trevithick-Sutton CC, Krogmann DW, Yeates TO (2003) The crystal structure of a cyanobacterial water-soluble carotenoid binding protein. Structure 11: 55–65

21. Konarev PV, Volkov VV, Sokolova AV, Koch MHJ, Svergun DI (2003) PRIMUS - a Windows-PC based system for small-angle scattering data analysis. J Appl Cryst 36: 1277–1282

22. Kumar S, Stecher G, Tamura K (2016) MEGA7: Molecular Evolutionary Genetics Analysis Version 7.0 for Bigger Datasets. Mol Biol Evol 33: 1870–1874

23. Lechno-Yossef S, Melnicki MR, Bao H, Montgomery BL, Kerfeld CA (2017) Synthetic OCP heterodimers are photoactive and recapitulate the fusion of two primitive carotenoproteins in the evolution of cyanobacterial photoprotection. Plant J 91: 646–656

24. Leverenz RL, Jallet D, Li MD, Mathies RA, Kirilovsky D, Kerfeld CA (2014) Structural and functional modularity of the orange carotenoid protein: distinct roles for the N- and C-terminal domains in cyanobacterial photoprotection. Plant Cell 26: 426–437

25. Leverenz RL, Sutter M, Wilson A, Gupta S, Thurotte A, de Carbon CB, Petzold CJ, Ralston C, Perreau F, Kirilovsky D, Kerfeld CA (2015) A 12 angstrom carotenoid translocation in a photoswitch associated with cyanobacterial photoprotection. Science 348: 1463–1466

26. Liu H, Zhang H, King j D, Wolf NR, Prado M, Gross ML, Blankenship RE (2014) Mass spectrometry footprinting reveals the structural rearrangements of cyanobacterial orange carotenoid protein upon light activation. Biochim Biophys Acta 1837: 1955–1963

27. Lopez-Igual R, Wilson A, Leverenz RL, Melnicki MR, Bourcier de Carbon C, Sutter M, Turmo A, Perreau F, Kerfeld CA, Kirilovsky D (2016) Different Functions of the Paralogs to the N-Terminal Domain of the Orange Carotenoid Protein in the Cyanobacterium Anabaena sp. PCC 7120. Plant Physiol 171: 1852–1866

28. Lu Y, Liu H, Saer R, Li VL, Zhang H, Shi L, Goodson C, Gross ML, Blankenship RE (2017) A Molecular Mechanism for Nonphotochemical Quenching in Cyanobacteria. Biochemistry 56: 2812–2823

29. Maksimov EG, Moldenhauer M, Shirshin EA, Parshina EA, Sluchanko NN, Klementiev KE, Tsoraev GV, Tavraz NN, Willoweit M, Schmitt FJ, Breitenbach J, Sandmann G, Paschenko VZ, Friedrich T, Rubin AB (2016) A comparative study of three signaling forms of the orange carotenoid protein. Photosynth Res 130: 389–401

30. Maksimov EG, Shirshin EA, Sluchanko NN, Zlenko DV, Parshina EY, Tsoraev GV, Klementiev KE, Budylin GS, Schmitt FJ, Friedrich T, Fadeev VV, Paschenko VZ, Rubin AB (2015) The Signaling State of Orange Carotenoid Protein. Biophys J 109: 595–607

31. Maksimov EG, Sluchanko NN, Mironov KS, Shirshin EA, Klementiev KE, Tsoraev GV, Moldenhauer M, Friedrich T, Los DA, Allakhverdiev SI, Paschenko VZ, Rubin AB (2017a) Fluorescent Labeling Preserving OCP Photoactivity Reveals Its Reorganization during the Photocycle. Biophys J 112: 46–56

32. Maksimov EG, Sluchanko NN, Slonimskiy YB, Mironov KS, Klementiev KE, Moldenhauer M, Friedrich T, Los DA, Paschenko VZ, Rubin AB (2017b) The Unique Protein-to-Protein Carotenoid Transfer Mechanism. Biophys J 113: 402–414

33. Maksimov EG, Sluchanko NN, Slonimskiy YB, Slutskaya EA, Stepanov AV, Argentova-Stevens AM, Shirshin EA, Tsoraev GV, Klementiev KE, Slatinskaya OV, Lukashev EP, Friedrich T, Paschenko VZ, Rubin AB (2017c) The photocycle of orange carotenoid protein conceals distinct intermediates and asynchronous changes in the carotenoid and protein components. Sci Rep 7: 15548

34. Melnicki MR, Leverenz RL, Sutter M, Lopez-Igual R, Wilson A, Pawlowski EG, Perreau F, Kirilovsky D, Kerfeld CA (2016) Structure, Diversity, and Evolution of a New Family of Soluble Carotenoid-Binding Proteins in Cyanobacteria. Molecular plant 9: 1379–1394

35. Moldenhauer M, Sluchanko NN, Buhrke D, Zlenko DV, Tavraz NN, Schmitt FJ, Hildebrandt P, Maksimov EG, Friedrich T (2017a) Assembly of photoactive orange carotenoid protein from its domains unravels a carotenoid shuttle mechanism. Photosynth Res 133: 327–341

36. Moldenhauer M, Sluchanko NN, Tavraz NN, Junghans C, Buhrke D, Willoweit M, Chiappisi L, Schmitt FJ, Vukojevic V, Shirshin EA, Ponomarev VY, Paschenko VZ, Gradzielski M, Maksimov EG, Friedrich T (2017b) Interaction of the signaling state analog and the apoprotein form of the orange carotenoid protein with the fluorescence recovery protein. Photosynth Res

37. Muzzopappa F, Wilson A, Yogarajah V, Cot S, Perreau F, Montigny C, Bourcier de Carbon C, Kirilovsky D (2017) Paralogs of the C-Terminal Domain of the Cyanobacterial Orange Carotenoid Protein Are Carotenoid Donors to Helical Carotenoid Proteins. Plant Physiol 175: 1283–1303

38. Panjkovich A, Svergun DI (2017) CHROMIXS: automatic and interactive analysis of chromatography-coupled small angle X-ray scattering data. Bioinformatics: btx846-btx846

39. Pascal AA, Liu Z, Broess K, van Oort B, van Amerongen H, Wang C, Horton P, Robert B, Chang W, Ruban A (2005) Molecular basis of photoprotection and control of photosynthetic light-harvesting. Nature 436: 134–137

40. Peschek GA, Obinger, Christian, Renger, Gernot (2011) Bioenergetic Processes of Cyanobacteria, 1 edn. Netherlands: Springer Netherlands.

41. Petoukhov MV, Franke D, Shkumatov AV, Tria G, Kikhney AG, Gajda M, Gorba C, Mertens HDT, Konarev PV, Svergun DI (2012) New developments in the ATSAS program package for small-angle scattering data analysis. J Appl Cryst 45: 342–350

42. Petoukhov MV, Svergun DI (2005) Global rigid body modeling of macromolecular complexes against small-angle scattering data. Biophys J 89: 1237–1250

43. Rambo RP, Tainer JA (2013) Accurate assessment of mass, models and resolution by small-angle scattering. Nature 496: 477–481

44. Robert X, Gouet P (2014) Deciphering key features in protein structures with the new ENDscript server. Nucleic Acids Res 42: W320–324

45. Shirshin EA, Nikonova EE, Kuzminov FI, Sluchanko NN, Elanskaya IV, Gorbunov MY, Fadeev VV, Friedrich T, Maksimov EG (2017) Biophysical modeling of in vitro and in vivo processes underlying regulated photoprotective mechanism in cyanobacteria. Photosynth Res 133: 261–271

46. Sluchanko NN, Klementiev KE, Shirshin EA, Tsoraev GV, Friedrich T, Maksimov EG (2017a) The purple Trp288Ala mutant of Synechocystis OCP persistently quenches phycobilisome fluorescence and tightly interacts with FRP. Biochim Biophys Acta 1858: 1–11

47. Sluchanko NN, Slonimskiy YB, Maksimov EG (2017b) Features of Protein—Protein Interactions in the Cyanobacterial Photoprotection Mechanism. Biochemistry (Moscow) 82: 1592–1614

48. Sluchanko NN, Slonimskiy YB, Moldenhauer M, Friedrich T, Maksimov EG (2017c) Deletion of the short N-terminal extension in OCP reveals the main site for FRP binding. FEBS Lett 591: 1667–1676

49. Sutter M, Wilson A, Leverenz RL, Lopez-Igual R, Thurotte A, Salmeen AE, Kirilovsky D, Kerfeld CA (2013) Crystal structure of the FRP and identification of the active site for modulation of OCP-mediated photoprotection in cyanobacteria. Proc Natl Acad Sci U S A 110: 10022–10027

50. Svergun DI (1992) Determination of the regularization parameter in indirect-transform methods using perceptual criteria. J Appl Cryst 25: 495–503

51. Svergun DI, Barberato C, Koch MHJ (1995) CRYSOL - a Program to Evaluate X-ray Solution Scattering of Biological Macromolecules from Atomic Coordinates. J Appl Cryst 28: 768–773

52. Svergun DI, Petoukhov MV, Koch MH (2001) Determination of domain structure of proteins from X-ray solution scattering. Biophys J 80: 2946–2953

53. Thurotte A, Bourcier de Carbon C, Wilson A, Talbot L, Cot S, Lopez-Igual R, Kirilovsky D (2017) The cyanobacterial Fluorescence Recovery Protein has two distinct activities: Orange Carotenoid Protein amino acids involved in FRP interaction. Biochim Biophys Acta 1858: 308–317

54. Thurotte A, Lopez-Igual R, Wilson A, Comolet L, Bourcier de Carbon C, Xiao F, Kirilovsky D (2015) Regulation of Orange Carotenoid Protein Activity in Cyanobacterial Photoprotection. Plant Physiol 169: 737–747

55. Valentini E, Kikhney AG, Previtali G, Jeffries CM, Svergun DI (2015) SASBDB, a repository for biological small-angle scattering data. Nucleic Acids Res 43: D357–363

56. Wilson A, Ajlani G, Verbavatz JM, Vass I, Kerfeld CA, Kirilovsky D (2006) A soluble carotenoid protein involved in phycobilisome-related energy dissipation in cyanobacteria. Plant Cell 18: 992–1007

57. Wilson A, Gwizdala M, Mezzetti A, Alexandre M, Kerfeld CA, Kirilovsky D (2012) The essential role of the N-terminal domain of the orange carotenoid protein in cyanobacterial photoprotection: importance of a positive charge for phycobilisome binding. Plant Cell 24: 1972–1983

58. Wilson A, Punginelli C, Couturier M, Perreau F, Kirilovsky D (2011) Essential role of two tyrosines and two tryptophans on the photoprotection activity of the Orange Carotenoid Protein. Biochim Biophys Acta 1807: 293–301

59. Wilson A, Punginelli C, Gall A, Bonetti C, Alexandre M, Routaboul JM, Kerfeld CA, van Grondelle R, Robert B, Kennis JT, Kirilovsky D (2008) A photoactive carotenoid protein acting as light intensity sensor. Proc Natl Acad Sci U S A 105: 12075–12080

60. Wu YP, Krogmann DW (1997) The orange carotenoid protein of Synechocystis PCC 6803. Biochim Biophys Acta 1322: 1–7

61. Yang J, Yan R, Roy A, Xu D, Poisson J, Zhang Y (2015) The I-TASSER Suite: protein structure and function prediction. Nat Methods 12: 7–8

